# Systematic discovery and perturbation of regulatory genes in human T cells reveals the architecture of immune networks

**DOI:** 10.1101/2021.04.18.440363

**Authors:** Jacob W. Freimer, Oren Shaked, Sahin Naqvi, Nasa Sinnott-Armstrong, Arwa Kathiria, Amy F. Chen, Jessica T. Cortez, William J. Greenleaf, Jonathan K. Pritchard, Alexander Marson

## Abstract

Complex gene regulatory networks ensure that important genes are expressed at precise levels. When gene expression is sufficiently perturbed it can lead to disease. To understand how gene expression disruptions percolate through a network, we must first map connections between regulatory genes and their downstream targets. However, we lack comprehensive knowledge of the upstream regulators of most genes. Here we developed an approach for systematic discovery of upstream regulators of critical immune factors – IL2RA, IL-2, and CTLA4 – in primary human T cells. Then, we mapped the network of these regulators’ target genes and enhancers using CRISPR perturbations, RNA-Seq, and ATAC-Seq. These regulators form densely interconnected networks with extensive feedback loops. Furthermore, this network is highly enriched for immune-associated disease variants and genes. These results provide insight into how immune-associated disease genes are regulated in T cells and broader principles about the structure of human gene regulatory networks.

**Highlights:**

- A systematic approach to identify upstream regulators of key immune genes in primary human cells
- Comprehensive RNA-Seq and ATAC-Seq perturbation maps after KO of individual discovered regulators
- Analysis uncovers a highly interconnected regulatory network of enhancers and genes in T cells
- This network is highly enriched for immune disease variants and genes shedding light on the trans-regulatory connections among key immune genes in health and disease

## Introduction

Human genetic studies have the potential to reveal underlying gene pathways that are disrupted to cause disease and point towards new targets for therapies. Over the past decade, genome wide association studies (GWAS) have identified thousands of disease-associated genetic variants (Buniello et al., 2019). However, it has been difficult to determine the functional consequences of these variants. Initial efforts have focused on mapping the cis-regulatory effects of genetic variants (Gallagher and Chen-Plotkin, 2018). However, in many cases, identifying the genes that are altered in cis does not provide a clear picture of disease etiology. Many of these cis-regulated genes are likely not directly involved in a relevant disease process, but instead trans-regulate genes that are directly involved (Liu et al., 2019; Võsa et al., 2018; Westra et al., 2013). Therefore, to identify the most salient disease genes and processes, we must map the genes that are affected in trans by disease-associated genetic variants. Currently, measuring trans-regulatory effects is proving harder than measuring cis-regulatory effects and is a largely unsolved problem.

There are only a few cases where the trans-regulatory impact of disease associated variants has been elucidated. An obesity-associated single nucleotide polymorphism (SNP) in the FTO locus alters the expression of IRX3 and IRX5, and in turn mitochondrial and thermogenesis genes in preadipocytes (Claussnitzer et al., 2015; Smemo et al., 2014). In another case, a SNP associated with type 2 diabetes is both a cis-eQTL (expression quantitative trait locus) for KLF14 and a trans-eQTL for a number of metabolism genes that are predicted KLF14 targets in adipose tissue (Small et al., 2018). In these cases, identifying the set of downstream genes affected by these SNPs enabled the authors to understand how each variant affects obesity or diabetes risk. However, mapping such trans-regulatory connections in human cells has been extremely challenging, making it difficult to understand how most disease associated variants alter disease relevant processes.

Mapping trans-eQTLs is difficult because they generally have very small effect sizes and therefore need very large sample sizes to detect (The GTEx Consortium, 2020; Võsa et al., 2018). An alternative approach is to experimentally perturb a gene and measure the resulting changes in expression of other genes. These regulatory connections are likely cell type-specific so must be mapped specifically in disease-relevant cell types (Califano et al., 2012). In order to tackle this problem, we have pioneered the use of CRISPR editing in freshly collected primary human T cells (Schumann et al., 2015). We recently developed a technology called SLICE (sgRNA Lentiviral Infection with Cas9 Electroporation) to perform large-scale CRISPR loss-of-function screening directly in primary human T cells (Shifrut et al., 2018). These approaches enable mechanistic studies directly in a cell type that plays critical roles in immune-mediated diseases.

Recent CRISPR-based methods such as Perturb-seq and CROP-seq have been widely adopted to knock out (KO) a selected set of genes and measure the resulting gene expression changes (Adamson et al., 2016; Datlinger et al., 2017; Dixit et al., 2016; Jaitin et al., 2016). We refer to this approach as “downstream mapping,” as it identifies genes that are downstream of the knocked-out genes in a transcriptional network. We have begun to pursue downstream mapping in primary human T cells (Schumann et al., 2020; Shifrut et al., 2018). However, such methods require a priori knowledge of the regulatory genes to select for disruption. In contrast, “upstream mapping” would enable us to start with genes of interest and unbiasedly discover the upstream regulators that control their expression. We could then draw regulatory connections between known disease genes and those upstream regulatory genes. Mapping these connections enables us to infer how disease-associated genetic variants that affect these upstream genes in cis, likely also affect these downstream disease genes in trans. However, upstream mapping has not been performed systematically in primary human cells.

Here, we coupled SLICE with fluorescence activated cell sorting (FACS) to identify the upstream regulators of the key immune genes: IL2RA (also known as CD25), IL-2, and CTLA4, in primary human CD4+ T cells. IL-2 is an important cytokine that can bind to the high affinity IL-2 receptor, IL2RA, to promote T cell proliferation and survival (Abbas et al., 2018; Spolski et al., 2018). CTLA4 limits T cells activation by inhibiting CD28-costimulation from CD80/CD86 on antigen presenting cells (Bayry, 2009). We focused on mapping the regulatory network around these three genes as the proper expression of these genes is critical for immune homeostasis and their disruption is associated with numerous complex and Mendelian immune diseases (Bezrodnik et al., 2014; Bousfiha et al., 2020; Caudy et al., 2007; Goudy et al., 2013; Kuehn et al., 2014; Linker-Israeli et al., 1983; Ochoa et al., 2021; Schubert et al., 2014; Sharfe et al., 1997).

After identifying upstream regulators, we performed downstream mapping to understand how these regulators affect other genes besides IL2RA, IL-2, and CTLA4. We individually knocked out 24 of the identified regulators and performed ATAC-Seq and RNA-Seq to directly measure changes in chromatin accessibility and gene expression. Combining both upstream and downstream mapping enabled us to generate the most comprehensive map of trans-regulatory connections in primary human cells (Figure 1A). Furthermore, these data provided novel insights into the regulatory architecture of human gene networks, revealing a highly interconnected network that contains extensive feedback loops, and is highly enriched for immune disease variants and genes.

**Figure 1:**
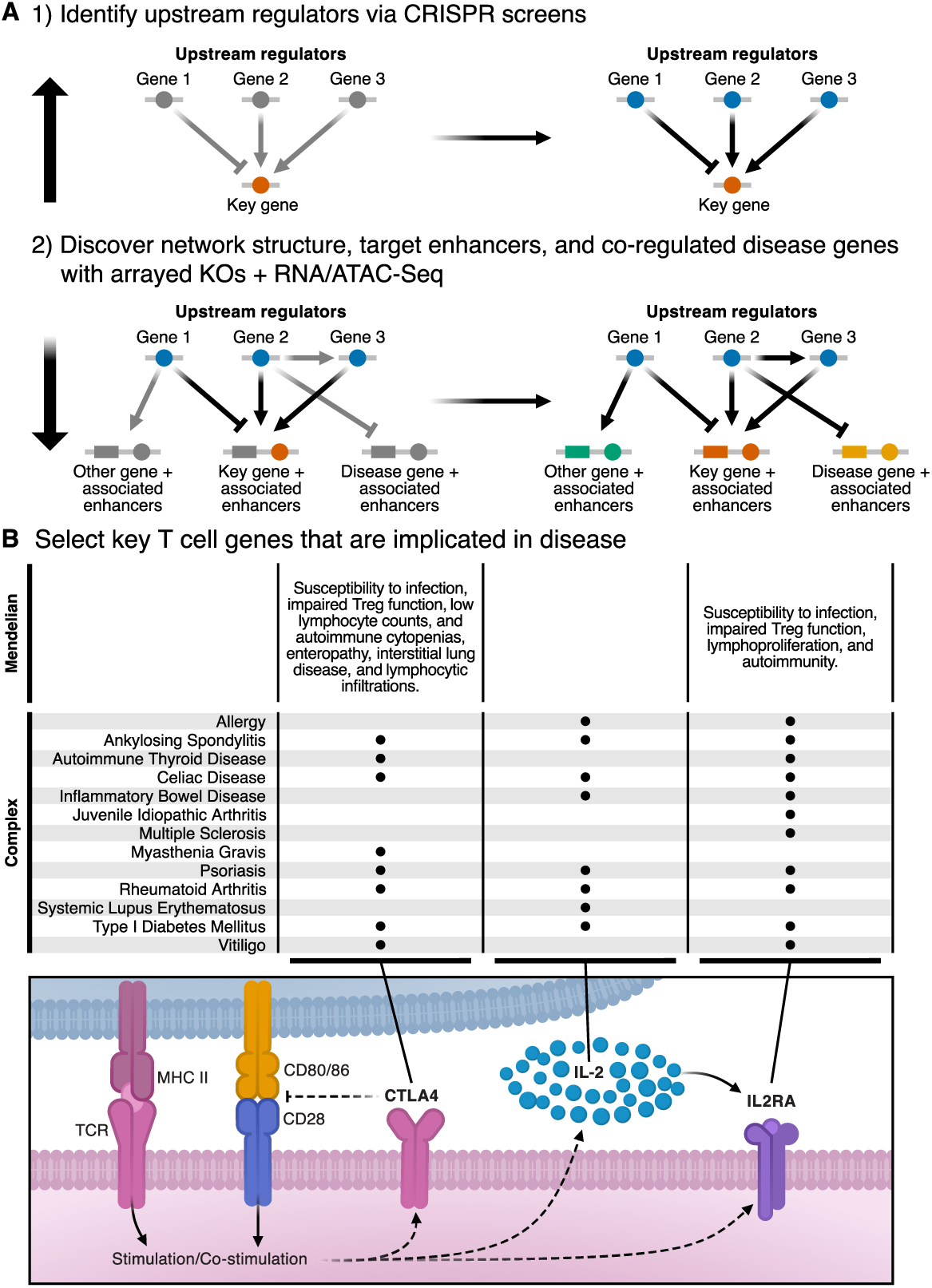
Approach to map disease gene networks in human T cells. **A)** 1) Perform pooled CRISPR loss-of-function screening coupled with fluorescence activated cell sorting to identify the upstream regulators of candidate key immune genes. 2) Individually knock out regulators identified in step one and measure downstream changes in chromatin accessibility and gene expression using ATAC-Seq and RNA-Seq to identify the network structure, target enhancers, and co-regulated disease genes. **B)** Schematic depicting the role of CTLA4, IL-2, and IL2RA in CD4+ T cells and a non-comprehensive list of common autoimmune diseases associated with dysregulation of these genes.

Our results provide a roadmap for identifying interconnected networks of disease associated genes by starting with a handful of impactful seed genes. We performed these experiments in CD4+ T cells as autoimmune disease associated SNPs are highly enriched in active chromatin regions in CD4+ T cells, making them an ideal target to study immune disease networks (Calderon et al., 2019; Farh et al., 2015; Finucane et al., 2018; Soskic et al., 2019). As we assemble more extensive maps of functional genetic networks in different human cell types, we can begin to identify and prioritize variants that disrupt important components of these networks and understand how each of these variants contributes to disease.

## Results

### Systematic discovery of upstream regulators of IL2RA, IL-2, and CTLA4 in primary human CD4+ T cells

As an initial step to understand the complete trans-regulatory wiring of human T cells, we first sought to measure a fraction of the global network centered around three important immune genes. Here we combined SLICE - to knock out thousands of genes in a pool of primary human T cells - with FACS to discover which factors act as upstream regulators of IL2RA, IL-2, and CTLA4 (Figure 2A). Since these three genes play critical roles in T cell biology - their dysregulation has been implicated in both complex and Mendelian forms of autoimmune disease - they constitute an ideal starting place to begin mapping immune gene regulatory networks (Figure 1B). We hypothesized that by building out the network around these key genes that we could identify central components of the immune regulatory network.

**Figure 2:**
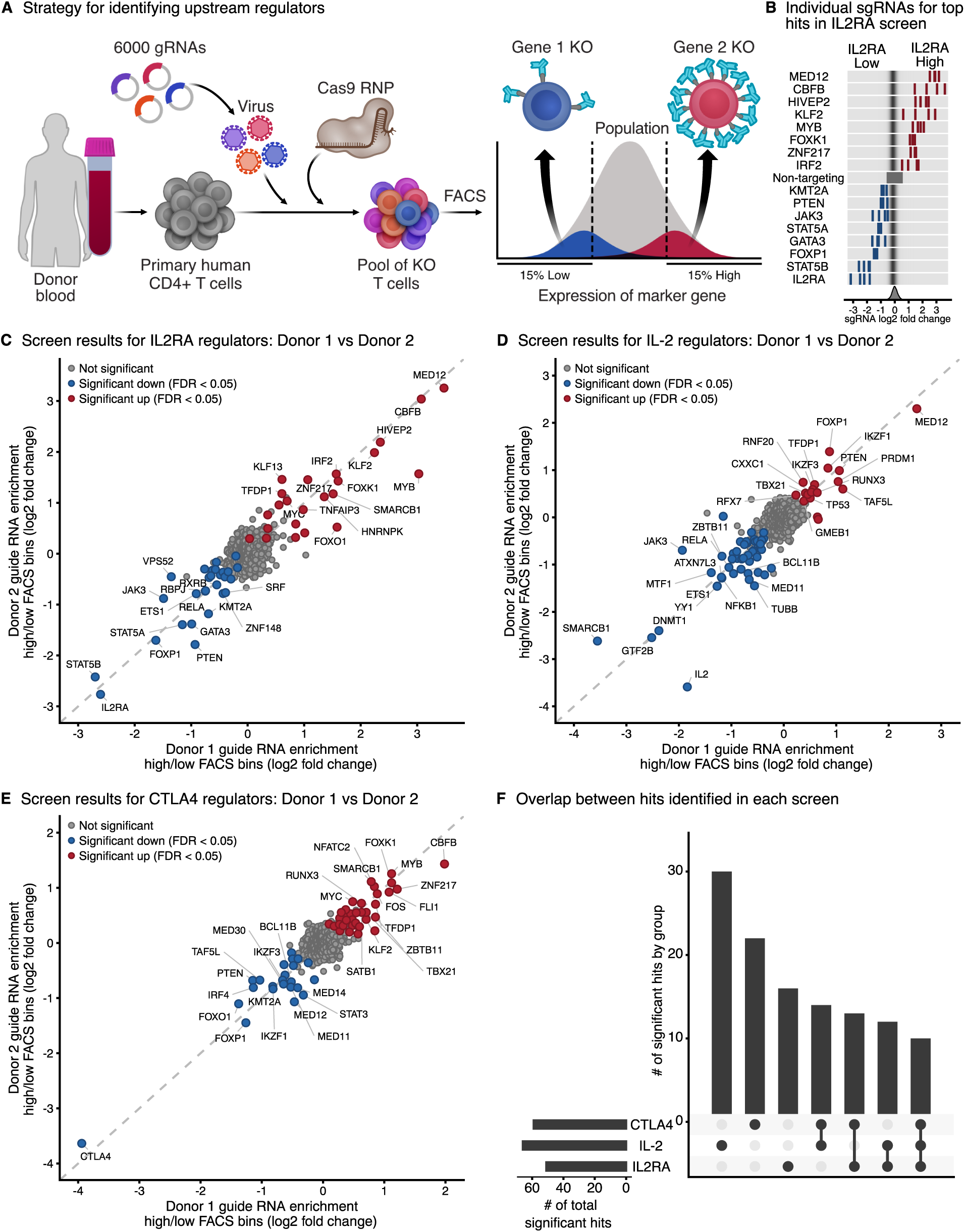
Systematic discovery of upstream regulators of IL2RA, IL-2, and CTLA4 in primary human CD4+ T cells. **A)** Strategy for identifying upstream regulators. We used SLICE (sgRNA Lentiviral Infection with Cas9 Electroporation) to generate a pool of knockout (KO) primary human CD4+ T cells. KO T cells were sorted into 15% high- or low-expression bins with FACS (fluorescence activated cell sorting) based on the expression of IL2RA, IL-2, or CTLA4. The guide RNAs in each bin were sequenced to identify the genes that positively or negatively regulate the expression of IL2RA, IL-2, or CTLA4. **B)** (top) Enrichment of individual guide RNAs in either the high- or low-expression bins for the top hits in the IL2RA screen. (bottom) Distribution of enrichment for all guide RNAs. **C-E)** Enrichment of guide RNAs in either the high- or low-expression bins for the IL2RA (n = 3), IL-2 (n = 2), and CTLA4 (n = 2) screens. Individual guide RNAs against the same gene were collapsed to the gene level. Significant hits with an FDR adjusted p-value < 0.05 across all donors are highlighted. **F)** The number of significant hits shared between the three screens. See also Figure S2.

Since we were interested in identifying trans-regulatory genes, we built a 6,000 guide RNA CRISPR library targeting 1,198 human transcription factors that have a known or inferred binding motif plus selected other candidate regulatory genes and controls (Table S1) (Lambert et al., 2018). We also included hits from other recent screens as well as a handful of genes of interest that might serve as important regulators in T cells (Cortez et al., 2020). We isolated CD4+CD25-T cells from healthy human blood donors, stimulated the cells, infected with lentivirus containing the guide RNA library and a GFP reporter, and then electroporated Cas9 ribonucleoprotein (RNP) to generate a pool of KO T cells (see methods for optimized SLICE protocol used here). We stained for IL2RA, IL-2, and CTLA4 and then used FACS to sort the cells with either high or low expression of these three proteins (Figures 2A and S2A). We sequenced guide RNAs in the high and low bins to determine which guide RNAs are differentially enriched and thus identify the genes that positively and negatively regulate the expression of IL2RA, IL-2, and CTLA4.

To confirm functional effects of SLICE editing, we initially compared the frequency of guide RNAs in the starting plasmid library to the abundance of each guide in the transduced GFP+ T cells. Guide RNAs targeting either GFP or known essential genes were highly depleted in the GFP+ sorted population (Figures S2B and S2D). Although we were focused on discovering regulators of IL2RA, IL-2, and CTLA4, we also identified a number of genes important for T cell proliferation or survival even in the short six-day screen window by comparing the guide RNA abundances in the GFP+ sorted population to the distribution in the starting plasmid library (Figure S2E). Guide RNAs targeting the tumor suppressors PTEN and P53 (Wang et al., 2018) became overrepresented during the course of the screen, while guide RNAs targeting components of the IL-2 signaling pathway (JAK3, IL2RA, IL2RB, STAT5B), which is important for T cell fitness (Liao et al., 2013; Ross and Cantrell, 2018; Spolski et al., 2018), were depleted (Figure S2E). Taken together, these results confirmed that we were able to uncover important regulators of cell state with the pooled functional perturbations in these experiments.

We next sorted the pool of KO T cells based on the levels of IL2RA, IL-2 or CTLA4 and identified guide RNAs that were over-represented in either the high or low expressing cell populations for each target protein. These IL2RA, IL-2, and CTLA4 screens respectively identified 51, 66, and 59 significant hits that were differentially enriched in either the 15% low or the 15% high bins (Figures 2C-E; Tables S2 and S3). Significant hits were highly reproducible between different biological donors, suggesting that the CRISPR perturbations identified robust gene regulatory relationships. Furthermore, distinct guide RNAs targeting the same gene tended to have concordant effects (Figure 2B). As an additional quality control, we confirmed that the hits are expressed in CD4+ T cells (Figure S2C). As expected, positive control guide RNAs targeting IL2RA, IL-2, and CTLA4 were among the most highly enriched in the low FACS bin in their respective screens. JAK3/STAT5 signaling positively regulates IL2RA expression, and we successfully detected JAK3, STAT5A, and STAT5B as strong positive regulators of IL2RA (Figure 2C) (Li et al., 2017; Liao et al., 2013; Spolski et al., 2018). We also identified a number of novel hits in each screen. For example, to our knowledge MED12 has not been implicated in IL-2 signaling, but we identified it as a regulator of both IL2RA and IL-2 (Figures 2C and 2D). Together, these results provide the most comprehensive picture of how the expression of IL2RA, IL-2, and CTLA4 are regulated in human T cells.

Many of the hits were shared among the screens, suggesting that IL2RA, IL-2, and CTLA4 are highly co-regulated. Of the 117 unique hits identified among the three screens, 39 were identified as significant regulators in two of the screens, and 10 were identified as significant regulators in all three (Figure 2F). For genes that were identified as regulators in multiple screens, we further analyzed whether they had concordant effects on IL2RA, IL-2, and CTLA4 levels. IL2RA promotes the fitness and proliferation of cells while CTLA4 inhibits T cell activation (Abbas et al., 2018; Bayry, 2009). The majority of genes that regulate both IL2RA and CTLA4, regulate them in the same direction, suggesting that this network may help to balance the consequences of T cell activation (Figure S2F). However, there were several perturbations that push IL2RA and CTLA4 in opposite directions and may be key drivers in dictating whether the overarching immune response is activating or inhibitory. These could represent interesting targets to either strongly activate or limit T cell stimulation in clinical settings.

### Arrayed KOs validate and characterize screen results

To validate and further characterize how these screen hits regulate IL2RA, IL-2, or CTLA4 we performed arrayed KOs using synthetic CRISPR RNA (crRNA)/Cas9 RNPs in 96-well plates (Figure 3A). We selected 50 genes, 3 positive controls, and 4 non-targeting controls for validation experiments. Thirty-nine of these genes regulate at least two out of three of the target genes, while the remaining genes were selected because they had large effects in an individual screen. For each gene, we used two guide RNAs per target and three donors per guide. We genotyped each KO sample using amplicon sequencing (Leenay et al., 2019). In many cases, over 90% of the amplicons contained insertions or deletions demonstrating highly efficient gene KO (Figures 3C and S3C-D). These high-efficiency gene ablations allowed us to assess altered cell phenotypes resulting from each regulatory gene disruption. Five days after RNP delivery we stained the cells and performed high-throughput flow cytometry to measure how each KO affected the protein levels of IL2RA, IL-2, and CTLA4.

**Figure 3:**
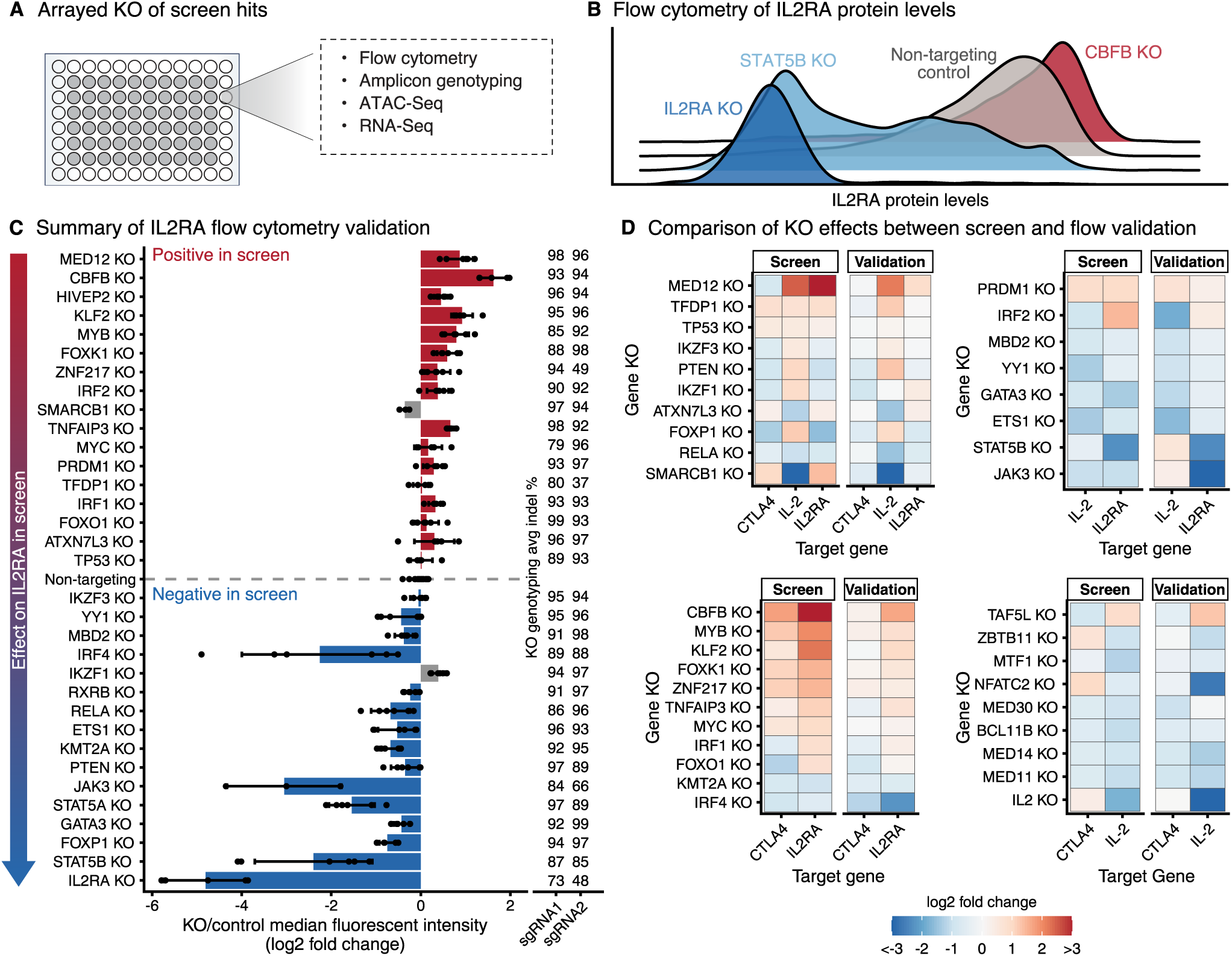
Arrayed KOs validate and characterize screen results. **A)** Arrayed KO of screen hits using CRISPR RNA (crRNA)/Cas9 ribonucleoproteins (RNPs) in 96-well plates, followed by in-depth phenotyping. **B)** Representative flow cytometry plots for IL2RA protein levels after KO of top screen hits. KOs that decrease or increase IL2RA levels are shown in blue or red respectively. **C)** Summary of changes in IL2RA levels measured using flow cytometry. Screen hits selected for validation are displayed on the Y-axis ordered by their effect size in the pooled CRISPR screen. For each KO, bars show the average change in IL2RA median fluorescence intensity relative to non-targeting controls. Dots show individual data points and error bars show standard deviation across two guide RNAs and three donors per guide RNA. Concordant changes between the screen and validation that increase or decrease IL2RA levels are shown in red or blue respectively. Discordant changes are shown in grey. The average insertion/deletion (indel) percentage at the genomic target site across multiple donors for guide RNA 1 (n = 3) and guide RNA 2 (n = 2) is shown to the right. **D)** Heatmaps summarizing the validation for IL2RA, IL-2, and CTLA4 screens focused on regulators that affect multiple targets. Heatmaps are grouped based on which of the three target genes a hit regulates. See also Figure S3.

Our pooled screens measured relative shifts in guide RNA frequency in the tails of the distribution of protein expression of IL2RA, IL-2, and CTLA4; the arrayed flow cytometry extended these measurements to show how each KO affected the full distribution of IL2RA, IL-2, and CTLA4 protein levels. Representative plots of IL2RA protein levels show that the KO of CBFP robustly increased IL2RA levels, while the KO of STAT5B robustly decreased IL2RA levels (Figure 3B). To summarize the results, we normalized median fluorescence intensity of each KO to the non-targeting controls on each plate. Overall, the results from the arrayed KOs were highly concordant with the results from the pooled CRISPR screens, confirming the biological reproducibility of our findings and demonstrating the power of the pooled screening approach (Figure 3C and S3A-D). We were particularly interested in the genes that co-regulate IL2RA, IL-2, and CTLA4, as these genes might be key regulators of important immune networks in T cells. The arrayed KO data were consistent with initial observations from the screens, confirming that key regulators control pairs of these key immune proteins and in some cases all three (Figure 3D). This validated dataset presents the most comprehensive set of regulatory connections between key immune genes and their upstream regulators in human T cells.

### Mapping genome-wide transcripts and chromatin sites controlled by each IL2RA regulator

Since many of the genes that we identified co-regulate IL2RA, IL-2, and CTLA4, we thought that our screen hits might control a broader, interconnected immune network. After successfully mapping the upstream regulators of IL2RA, IL-2, and CTLA4 we switched to downstream mapping to identify what other genes are in this network (Figure 1A). We focused on the IL2RA regulatory network and selected 24 regulators that had the largest effects on IL2RA levels in the validation dataset, including IL2RA itself. For a control comparison, we selected eight guide RNAs that target the safe harbor locus *AAVS1*. To identify each regulators’ downstream target genes and enhancers, we performed arrayed RNP KOs in CD4+ T cells from three human blood donors followed by bulk RNA-Seq and ATAC-Seq (Figure 3A; Tables S4, S5, S6). This approach enabled us to measure thousands of additional gene expression and chromatin changes compared to alternative single-cell sequencing methods.

We first confirmed that the RNA-Seq data revealed meaningful changes in the CRISPR KOs. Consistent with the genotyping results that showed high rates of insertion/deletion mutations at the target sites (Figure 3C), RNA-Seq demonstrated that the expression of most targeted regulators was decreased in their respective KO RNA-Seq samples (Figure S4A). The few examples where this was not the case could be due to a lack of nonsense-mediated decay and/or feedback mechanisms on transcription. Furthermore, flow cytometry measurements of IL2RA protein levels in the same RNP-targeted cell populations collected for RNA-Seq revealed that changes in IL2RA mRNA levels and IL2RA protein levels were highly correlated (Figure S4B). Although the readout for our screen was IL2RA protein levels, the regulators that we identified also affect *IL2RA* transcript levels. These results confirm that the regulatory effects of KOs can be ascertained from the RNA-Seq data.

CRISPR KO also affected the global chromatin landscape as assessed with ATAC-Seq. Sites of chromatin accessibility that were altered upon ablation of a specific transcription factor tended to be enriched for the corresponding binding sequence motif (see methods). In these KO samples, we analyzed how many differential ATAC-Seq peaks contained a motif associated with the ablated transcription factor. On average, 41% of differential ATAC-Seq peaks contained a matching motif, suggesting that we captured many primary regulatory effects (Figure S4C). These data reveal specific transcription factors required for maintaining chromatin accessibility in human CD4+ T cells at sites throughout the genome.

The number of accessible chromatin regions and genes affected by each KO varied greatly. For example, in the ATAC-Seq data, the loss of FOXK1 affected only 548 peaks, while the loss of CBFB affected 34,379 peaks (FDR adjusted p-value < 0.05, Figure S4E). In the RNA-Seq data, the loss of HIVEP2 significantly affected the expression of only 19 genes, while the loss of MED12 significantly affected the expression of 4,641 genes (FDR adjusted p-value < 0.05, Figure S4D). While the number of significant changes in gene expression and chromatin accessibility were roughly correlated, there were interesting exceptions: the loss of YY1 produces many changes in gene expression, but comparatively few changes in chromatin accessibility. These data provide a view of how broadly the chromatin landscape and transcriptome depends on each individual regulator identified by the screens.

### Individual regulators maintain chromatin accessibility at distinct enhancers to control IL2RA gene expression

We next analyzed how the loss of each IL2RA regulator affected gene expression and altered chromatin accessibility throughout the *IL2RA* locus. Overall, the loss of negative regulators of IL2RA not only increased *IL2RA* expression, but also tended to increase chromatin accessibility (Figure 4A). Conversely, the loss of positive regulators of IL2RA decreased *ILRA* expression and chromatin accessibility (Figure 4A). More interestingly, we observed discrete effects of different KOs on accessibility at different non-coding elements in the *IL2RA* locus (Figure 4B). The loss of both CBFB and TNFAIP3 both increased the expression of *IL2RA* and chromatin accessibility. However, loss of CBFB resulted in a broad domain of increased accessibility around an enhancer in the first intron of *IL2RA* that was unaffected by TNFAIP3 KO. The loss of TNFAIP3 increased accessibility at an enhancer 3’ of *IL2RA* that was largely unchanged in the CBFB KO. In contrast, the loss of both IRF4 and STAT5B decreased chromatin accessibility and the expression of *IL2RA*. The loss of IRF4 decreased accessibility at the 3’ enhancer, which was minimally changed in the STAT5B KO. On the other hand, the loss of STAT5B decreased accessibility at two intronic enhancers that were minimally changed in the IRF4 KO. These examples highlight how different transcriptional pathways act on distinct enhancer elements to precisely control the levels of IL2RA. Broadly these data demonstrate how genetic perturbations can be coupled with ATAC-Seq and RNA-Seq to build gene networks tying regulators to both specific enhancer elements and downstream target gene expression changes.

**Figure 4:**
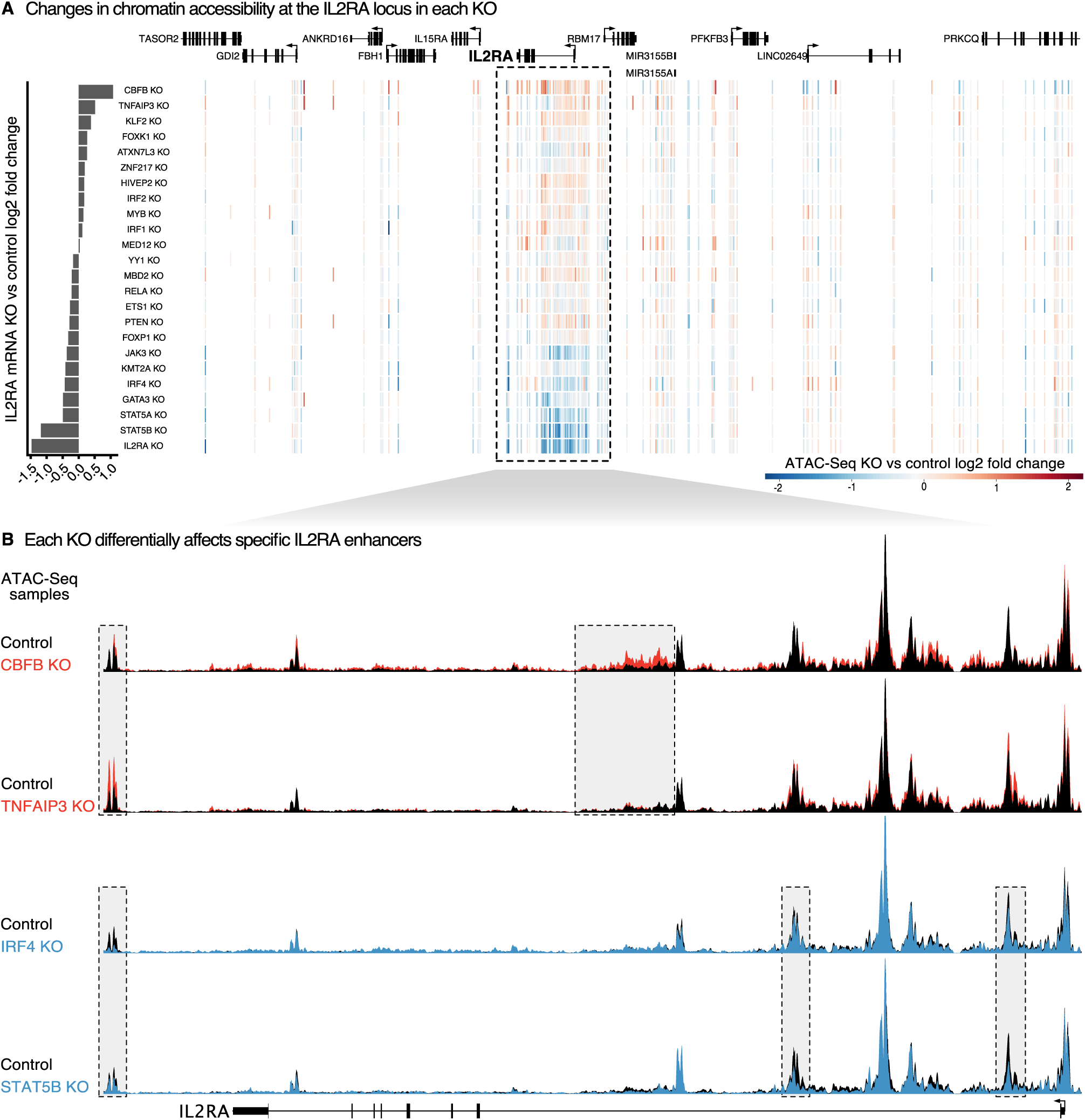
Individual regulators maintain chromatin accessibility at distinct enhancers to control IL2RA gene expression. **A)** Heatmap of changes in chromatin accessibility around the IL2RA locus. Each row represents a different KO and each vertical line is an ATAC-Seq peak. Peaks with increased or decreased accessibility are shown in red or blue respectively. The graph at the left shows the change in IL2RA mRNA expression in each KO. **B)** Changes in chromatin accessibility at specific enhancers throughout the IL2RA locus in CBFB, TNFAIP3, IRF4, and STAT5B KOs. Examples of enhancer dynamics that differ between the KO samples are boxed. See also Figure S4.

### IL2RA regulators form highly interconnected gene networks

Compared to bacteria and yeast, we know significantly less about the architecture of gene networks in human cells. Large perturbation studies in these single cell model organisms have placed genes in pathways and identified common regulatory features of these pathways such as feedback loops (Alon, 2007; Hughes and de Boer, 2013; Kemmeren et al., 2014). However, as it has been difficult to directly perturb primary human cells, efforts to construct human gene networks have largely relied on observational data such as analyzing gene co-expression data (Aibar et al., 2017; van Dam et al., 2018; Fiers et al., 2018; Langfelder and Horvath, 2008; Margolin et al., 2006). However, such observational data alone cannot determine the direction of effects in gene networks.

The knockout of 24 genes followed by RNA-Seq and ATAC-Seq is one of the largest datasets yet generated to understand how perturbations affect gene expression and chromatin regulation in primary human cells. As each of the genes that we disrupted affects IL2RA expression, we wondered if they act as part of a large, interconnected network or rather act independently. We first analyzed how each KO affects the expression of the other IL2RA regulators. The loss of each regulator significantly altered the expression of between 1 and 18 (median 9.5) out of 24 other IL2RA regulators (including IL2RA itself) (Figure 5A). The transcription factor IRF4 exemplifies the extensive regulatory connections observed among IL2RA regulators. KO of IRF4 significantly altered the expression of 9 other IL2RA regulators, while KO of 15 other IL2RA regulators significantly altered the expression of IRF4 (Figure 5B). The number of connections between IRF4 and other regulators illustrates how even this subnetwork of 24 genes is highly interconnected; perturbing any of the regulators in this subnetwork produces cascading effects on the other regulators and in turn their downstream targets. As we only performed RNA-Seq with 24 of our screen hits ablated, the full network is likely significantly more interconnected than previously appreciated.

**Figure 5:**
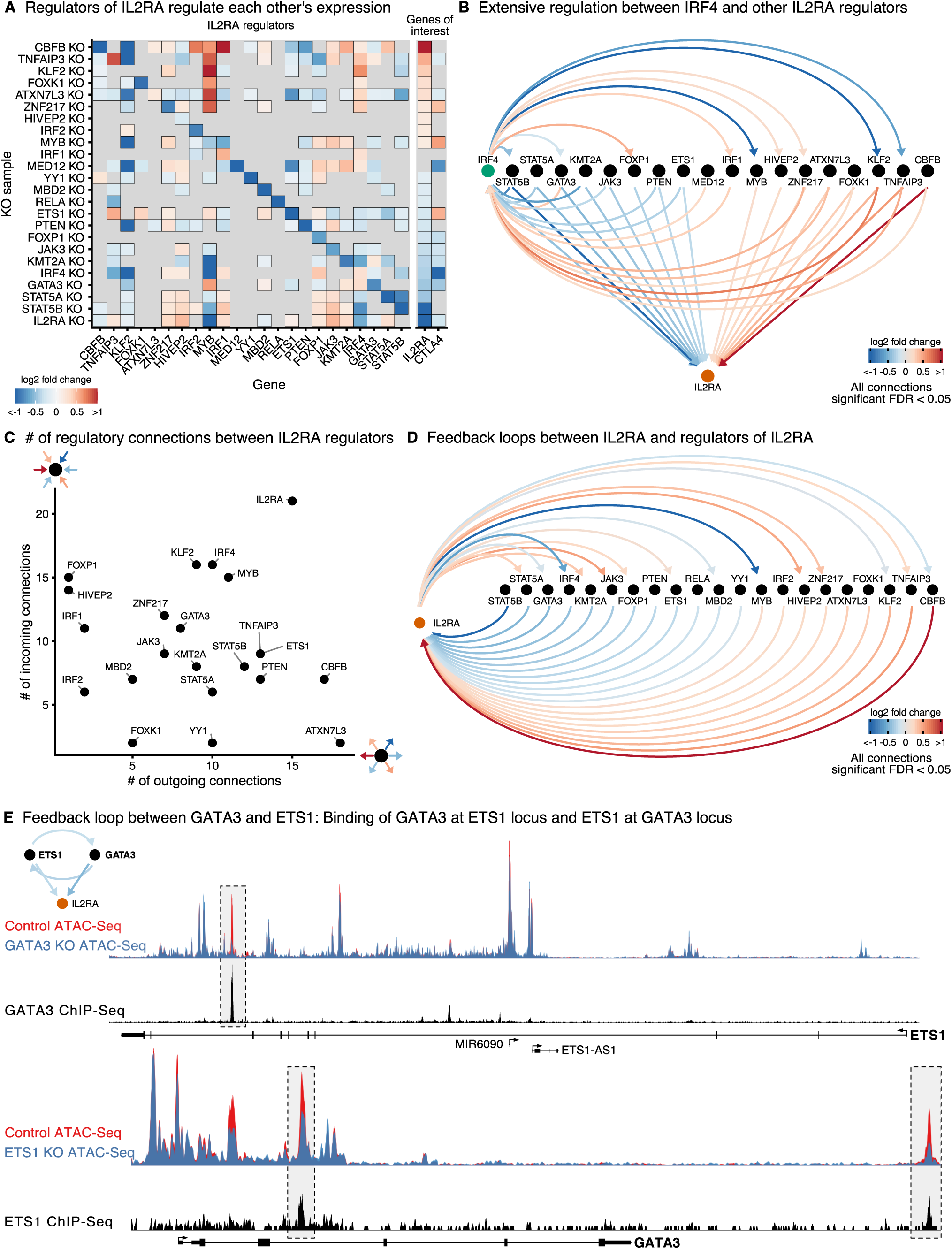
IL2RA regulators form highly interconnected gene networks. **A)** Significant changes in the expression of IL2RA regulators, IL2RA, and CTLA4 in each KO sample (FDR adjusted p-value < 0.05 for all changes shown). **B)** Map of regulatory connections between IRF4 and other IL2RA regulators detected via RNA-Seq in each KO sample. Arrows point towards the target gene perturbed in each KO sample, while the color of lines shows the fold change of the target (FDR adjusted p-value < 0.05 for all changes shown). **C)** The number of outgoing and incoming regulatory connections between each IL2RA regulator and all other IL2RA regulators. **D)** Map of regulatory connections between IL2RA and regulators of IL2RA as described in B. **E)** Feedback loop between ETS1 and GATA3; connections as described in B. (top tracks) Changes in chromatin accessibility at the ETS1 locus in GATA3 KO cells and ChIP-Seq of GATA3 binding at the ETS1 locus. (bottom tracks) Changes in chromatin accessibility at the GATA3 locus in ETS1 KO cells and ChIP-Seq of ETS1 binding at the GATA3 locus. (GATA3 ChIP-Seq from <u>(Kanhere et al., 2012)</u> and ETS1 ChIP-Seq from <u>(Schmidl et al., 2014)</u>).

In order to better understand the architecture of this network, we analyzed the number of “outgoing” and “incoming” connections for each IL2RA regulator (Figure 5C). These connections do not distinguish direct vs. indirect regulation, but represent cumulative causal changes upon the loss of each regulator. Outgoing connections represent the number of other IL2RA regulators affected by a given KO. Incoming connections represent the number of KOs that affect a given regulator. CBFB and ATXN7L3 have many outgoing connections, but only a few incoming connections suggesting that they are more broad regulators. Conversely, HIVEP2 and FOXP1 have many incoming connections, but very few outgoing connections revealing that they are relatively more targeted in regulating IL2RA.

Strikingly, IL2RA itself has a high number of both incoming and outgoing connections (Figure 5C). The high number of incoming connections for IL2RA was expected, given our experimental design, but the number of outgoing connections suggests that IL2RA is involved in extensive feedback loops. We analyzed how many IL2RA regulators were differentially expressed in the IL2RA KO data and observed numerous feedback loops between IL2RA regulators and IL2RA itself (Figure 5D). We also identified numerous feedback loops between IL2RA regulators. For example, the loss of ETS1 decreased *GATA3* expression while the loss of GATA3 decreased *ETS1* expression (Figure 5E). To determine if these transcription factors directly regulate each other’s expression, we analyzed public ETS1 and GATA3 ChIP-Seq in human CD4+ T cells along with our ETS1 and GATA3 KO ATAC-Seq data (Kanhere et al., 2012; Schmidl et al., 2014). In the GATA3 KO ATAC-Seq data, there was a specific loss of chromatin accessibility that overlaps a GATA3 ChIP-Seq peak at the *ETS1* locus. Similarly, in the ETS1 KO ATAC-Seq data, there was a loss of chromatin accessibility that overlaps two ETS1 ChIP-Seq peaks at the *GATA3* locus. Together, these data suggest that ETS1 and GATA3 directly control specific sites of chromatin accessibility at each other’s locus and regulate each other’s expression in human CD4+ T cells. To date, feedback loops have likely been under-detected in human gene networks because their identification requires the reciprocal perturbations performed in this study. Across our KOs, we discovered that feedback loops are a common feature of human gene networks. Additional future perturbation experiments in diverse primary cell types and conditions will continue to reveal the full complexity of human gene networks.

### Highly co-regulated gene sets are enriched for immune disease genes

Since the IL2RA regulators form a highly interconnected network, we were interested in whether they co-regulate a set of target genes with critical roles in T cell functions. We identified genes that are co-regulated to different degrees with IL2RA based on the number of KO samples where the gene is differentially expressed (Figure 6A). We hypothesized that the set of genes that depended on many of the same regulators as IL2RA may also be important for healthy T cell function. Most genes were differentially expressed in between one and six KOs, but hundreds of genes were differentially expressed in ten or more KOs indicating that they are highly co-regulated by the IL2RA regulators (Figure 6B). These highly co-regulated genes are significantly enriched for annotated immune genes (Figure 6B). We were interested in whether the IL2RA regulators also co-regulate immune disease-associated genes besides IL2RA, CTLA4, and IL-2. We identified differentially expressed Mendelian immune disease genes based on a curated list of inborn errors of immunity (IEI) genes (Tangye et al., 2020, 2021). We also identified differentially expressed genes that are within 100kb of autoimmune GWAS SNPs (Taylor et al., 2021). Strikingly, the IL2RA co-regulated network was significantly enriched for both these Mendelian and GWAS disease genes (Figure 6B). This finding suggests that key immune disease genes sit in a highly connected central network. This network structure is consistent with models suggesting that many peripheral regulatory connections converge on a set of core genes that are directly involved in disease relevant cellular processes (Boyle et al., 2017; Liu et al., 2019).

**Figure 6:**
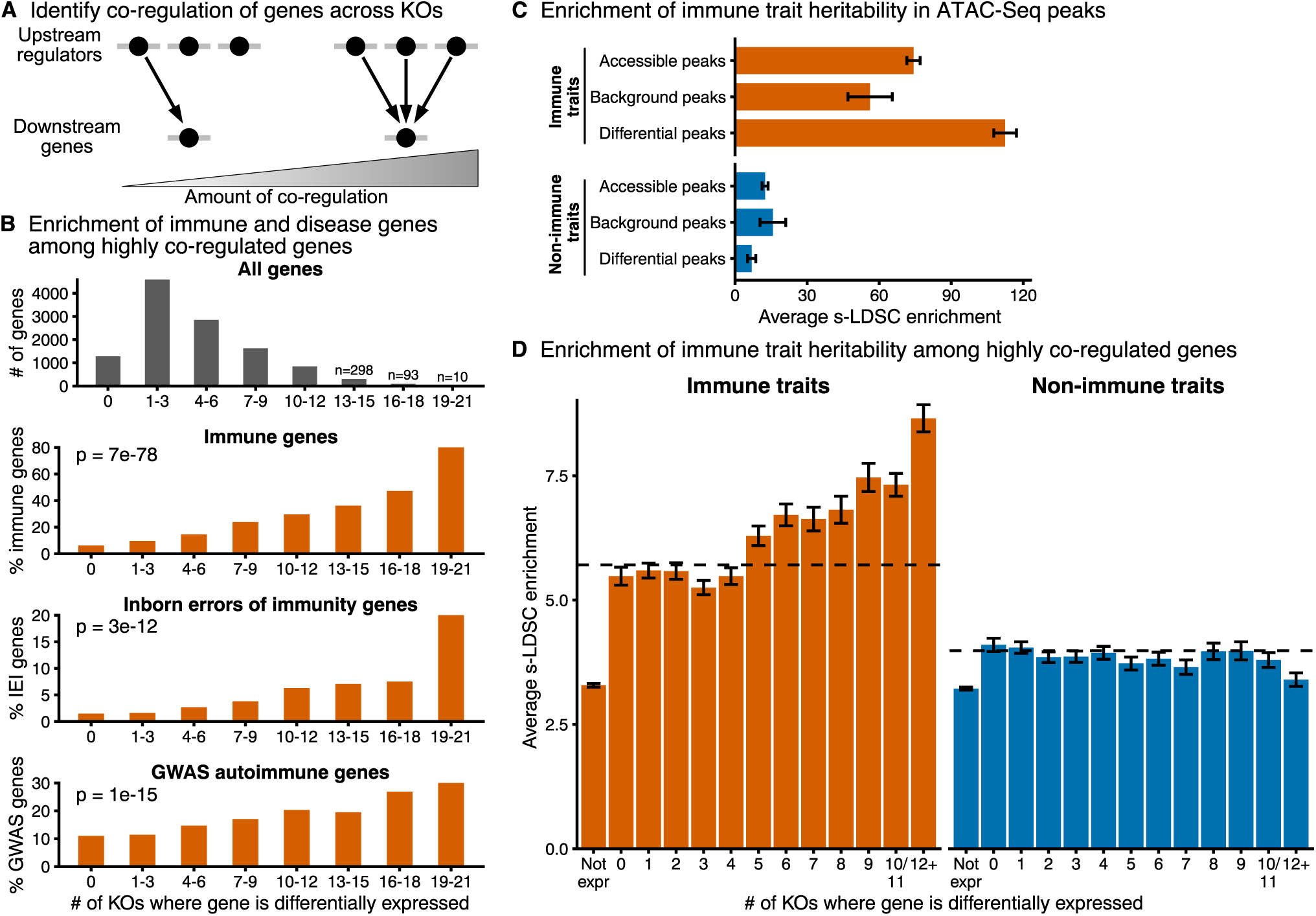
Highly co-regulated gene sets are enriched for immune disease genes. **A)** Define the degree of gene co-regulation based on the number of KO samples where the gene is differentially expressed. **B)** The total number of genes that are significantly differentially expressed in each KO bin or the percent of differentially expressed genes that are classified as “immune system process” genes by gene ontology, inborn error of immunity (IEI) Mendelian disease genes, or autoimmune GWAS genes. (P-values calculated with a logistic regression using average expression and co-regulation bins as inputs. IL2RA, IL-2, and CTLA4 excluded from analysis). **C-D)** Enrichment of heritability for immune traits compared to non-immune traits in all ATAC-Seq peaks or significant differentially accessible ATAC-Seq peaks (C) or in a window of 100kb around highly co-regulated genes (D). Average background enrichment across co-regulation bins is shown as dashed lines in (D). Enrichment calculated using stratified LD score regression. Traits were meta-analyzed using inverse-variance weighting; average enrichment and standard deviation shown. See also Figure S6.

To further explore how IL2RA regulators and their targets might be involved in complex forms of immune disease, we used stratified Linkage Disequilibrium Score (s-LDSC) regression to calculate the enrichment of immune and non-immune trait heritability in ATAC-Seq peaks (Finucane et al., 2015). Consistent with previous analyses, heritability for immune traits is highly enriched in T cell accessible chromatin compared to non-immune traits (Figure 6C) (Calderon et al., 2019; Farh et al., 2015; Finucane et al., 2018; Maurano et al., 2012; Soskic et al., 2019). We next asked if the loss of IL2RA regulators affected the accessibility of enhancers containing immune trait SNPs. ATAC-Seq peaks that were significantly changed in at least one KO sample were further enriched for immune trait heritability (Figure 6C). This enrichment suggests that IL2RA regulators control a set of critical chromatin accessibility sites that can be altered by genetic variants and contribute to immune disease risk, revealing a network of co-regulated non-coding elements that could help to prioritize and characterize candidate GWAS hits for various immune diseases.

We next wanted to test whether highly co-regulated genes are specifically enriched for immune trait heritability and not complex trait heritability more generally. We used s-LDSC to measure the enrichment of heritability in regions surrounding genes co-regulated to different degrees for a wide range of immune and non-immune traits (Finucane et al., 2018). Highly co-regulated genes were enriched for immune trait heritability compared to non-immune traits or immune genes that are not highly co-regulated (Figure 6D). To control for immune relevant genes being more highly expressed in T cells, and thus easier to detect as differentially expressed, we generated a background set of genes for each co-regulation bin. These background sets were sampled from genes that were differentially expressed in less than five samples and were matched to the differentially expressed genes in each co-regulation bin based on expression. Highly co-regulated genes were more enriched than their corresponding background sets, demonstrating that the enrichment is not just driven by levels of expression (Figures 6D and S6A). Overall, these data reveal that regulators of IL2RA also co-regulate the expression of a network of other Mendelian and complex immune disease genes which underlie multiple autoimmune and immune dysregulation diseases. Furthermore, this process of identifying disease networks by first identifying the regulators of key Mendelian disease genes could be replicated in other human cell types as a general approach to help map and understand disease variants and the complement of genes associated with different cell types function and dysregulation.

### IL2RA regulators affect enhancers and genes associated with multiple sclerosis

Among immune traits analyzed, multiple sclerosis (MS) heritability was enriched markedly in accessible ATAC-Seq peaks in our dataset (Figure S7A). To explore how IL2RA regulators affect enhancers and genes near SNPs associated with MS, we analyzed genome-wide significant SNPs reported in a recent meta-analysis of MS GWAS (International Multiple Sclerosis Genetics Consortium, 2019). To identify the likely causal SNP among GWAS hits, we previously developed an algorithm called Probabilistic Identification of Causal (PICS) (Farh et al., 2015; Taylor et al., 2021). There are 100 loci that contain a single MS-associated SNP with a PICS probability greater than 50% (Figure 7A). Twenty-eight of these SNPs are within an open chromatin ATAC-Seq peak in our CD4+ T cell data. Remarkably, 17 of these SNPs fall within an ATAC-Seq peak that was altered by loss of one or more IL2RA regulators (Figure 7A; p-value = 0.004 hypergeometric test). This reveals specific regulators that are required to regulate non-coding elements that harbor fine-mapped MS variants and places a large set of MS variants into a newly defined gene regulatory network.

**Figure 7:**
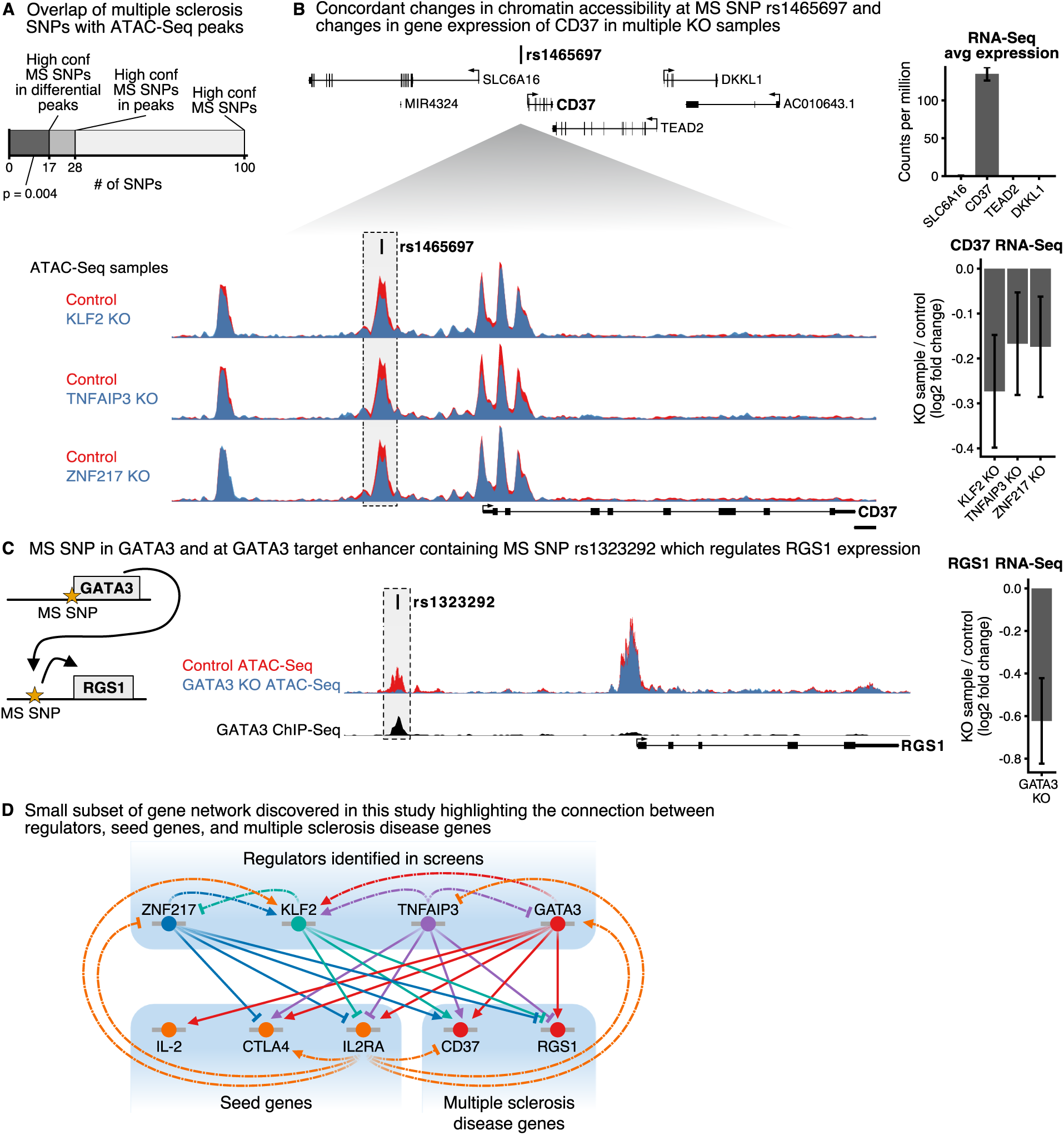
IL2RA regulators affect enhancers and genes associated with multiple sclerosis. **A)** The number of high confidence multiple sclerosis (MS) SNPs with a PICS probability > 0.5 in the genome, in all ATAC-Seq peaks, or in differentially accessible ATAC-Seq peaks (for which the p-value was calculated with hypergeometric test). **B)** (top) The expression of genes surrounding MS SNP rs1465697 in CD4+ T cells. (bottom) Changes in chromatin accessibility at rs1465697 and accompanying changes in CD37 mRNA expression in KLF2, TNFAIP3, and ZNF217 KOs. **C)** Cartoon illustrating MS SNPs both at GATA3 and at an enhancer upstream of RGS1. Changes in chromatin accessibility at rs1323292 and accompanying changes in RGS1 mRNA expression in GATA3 KO. ChIP-Seq of GATA3 binding at rs1323292. **D)** A small subset of the regulatory connections identified in this study between seed immune disease genes, their upstream regulators, and MS disease genes. This subnetwork is focused just on regulators that have a significant effect on CD37 or RGS1 chromatin accessibility and gene expression. See also Figure S7.

To explore further how disease SNPs might disrupt a regulatory cascade within a gene network, we sought to link KO of IL2RA regulators to changes in enhancers containing MS SNPs and to changes in the expression of nearby genes. For each SNP within a differential ATAC-Seq peak, we analyzed how many protein coding genes within 100kb are differentially expressed in at least one KO sample (Figure S7B). The MS SNP rs1465697 has been reported as an eQTL for CD37, DKKL1, TEAD2, and SLC6A16 (Ochoa et al., 2021). However, of the seven genes within 100kb of rs1465697, only *CD37* was expressed in our T cell RNA-Seq data (Figure 7B). The loss of KLF2, TNFAIP3, and ZNF217 all decreased chromatin accessibility at the enhancer containing rs1465697 and decreased the expression of *CD37* (Figure 7B). The concordant changes in chromatin accessibility and gene expression across different KOs strongly link this enhancer to CD37 as the relevant target gene. Together, these results suggest that rs1465697 might alter the binding of transcription factors within this enhancer, which depends on KLF2, TNFAIP3, and ZNF27 for accessibility, and disrupt the normal expression of *CD37*. Analyzing concordant effects on specific enhancers and nearby genes across multiple KOs could be a generalizable strategy to functionally link enhancers to their target genes.

Distinct SNPs associated with the same disease might also disrupt multiple components of a critical regulatory cascade. There is an MS SNP within the first intron of the gene encoding the transcription factor GATA3, suggesting that altered GATA3 levels contribute to MS risk. We asked whether any high confidence MS SNPs are located in enhancers that are differentially accessible upon KO of GATA3. Using public GATA3 ChIP-Seq in CD4+ T cells (Kanhere et al., 2012), we found a putative enhancer directly bound by GATA3 upstream of *RGS1* that harbors the MS SNP rs1323292 (Figure 7C). Furthermore, KO of GATA3 significantly decreased chromatin accessibility at this enhancer and the expression of *RGS1* (Figure 7C). These data suggest that RGS1 and its MS-associated enhancer directly depend on GATA3. Furthermore, these data demonstrate how disease SNPs might affect multiple genes within a single regulatory cascade. Together, these examples illustrate how we can combine genetic perturbations, ATAC-Seq, and RNA-Seq in order to understand how disease SNPs disrupt regulatory connections within gene networks, highlighting key regulators, enhancers, and downstream target genes required for immune homeostasis (Figure 7D).

## Discussion

Despite the recognition that we must understand how genetic variants disrupt trans-regulatory networks to understand how they contribute to disease risk, there are only a handful of examples where it has been possible to identify the relevant trans-regulated targets of a disease associated variant (Claussnitzer et al., 2015; Small et al., 2018; Smemo et al., 2014). To map trans-regulatory networks in humans, the field has traditionally faced a tradeoff between performing observational studies in disease relevant primary cells or functional studies in less relevant cell lines or animal models. To overcome limitations with each of these approaches, we combined CRISPR perturbations, RNA-Seq, and ATAC-Seq in primary human T cells to directly measure the regulation of human disease genes and enhancers.

Regulatory connections between genes in primary human cells can be inferred through trans-eQTL studies which measure associations between gene expression levels and genetic variation across individuals. However, trans-eQTL effects are small and require large sample sizes to detect. The latest version of GTEx detected 4,278,636 cis-eQTLs, but only 143 trans-eQTLs (The GTEx Consortium, 2020). A much larger meta-analysis across 31,684 individuals identified that 3,853 of 10,317 trait-associated SNPs are trans-eQTLs (Võsa et al., 2018). These studies demonstrate that using trans-eQTLs to identify trans-regulatory connections will require very large sample sizes. Furthermore, such studies will have to be repeated across a large number of individuals in each cell type of interest. In contrast, here we identified dozens of upstream regulators of IL2RA, IL-2, and CTLA4 and identified thousands of downstream targets for 24 IL2RA regulators by performing genetic perturbations directly in disease relevant primary cells.

Functional studies to identify upstream regulators have primarily been performed in human cell lines or in mouse models, which are not ideal for understanding disease biology. Brockmann et al. used random mutagenesis in a human haploid cell line to identify the regulators of proteins involved in diverse cell processes (Brockmann et al., 2017). Our lab and others have combined CRISPR perturbations in mouse immune cells with FACS to identify genes that regulate the induction of TNF in response to LPS, Th2 differentiation, or the stability of regulatory T cells marked by the expression of Foxp3 (Cortez et al., 2020; Henriksson et al., 2019; Parnas et al., 2015). Here, we extended these studies to primary human cells to map disease relevant gene networks. We identified regulators that not only control the expression of the disease genes IL2RA, IL-2, and CTLA4, but also regulate a network of other immune disease genes and enhancers. We demonstrated that a subset of these regulators control enhancers containing MS SNPs and linked these enhancers to their credible target genes. Using primary human cells enabled us to integrate the regulatory connections that we identified with existing human genetics data to gain insight into the regulatory effects of immune disease variants.

We were able to build comprehensive regulatory maps between 24 IL2RA regulators and thousands of downstream target genes and enhancers. While gene co-expression data is commonly used to construct human gene regulatory networks, such observations are merely correlative (Aibar et al., 2017; van Dam et al., 2018; Fiers et al., 2018; Langfelder and Horvath, 2008; Margolin et al., 2006). However, previous efforts to map gene networks using perturbations were limited by the number of genes studied, less robust siRNA perturbations, or less sensitive microarray measurements (Amit et al., 2009; Cusanovich et al., 2014). More recently, a number of studies have coupled CRISPR perturbations with single cell sequencing to map regulatory networks. Consistent with our observations, several of these studies observed highly interconnected regulation between a limited set of transcription factors (Dixit et al., 2016; Qiu et al., 2020; Rubin et al., 2019). However, since we used Cas9 RNPs to generate highly pure KO populations, we were able to perform bulk ATAC-Seq and RNA-Seq to capture thousands of additional chromatin and gene expression changes compared to single cell sequencing. Bulk profiling enabled us to build much more comprehensive regulatory maps, especially between transcription factors which tend to be lowly expressed and thus can be more difficult to measure using single cell RNA-Seq (Vaquerizas et al., 2009).

These regulatory maps also revealed the highly interconnected structure of human gene networks. The IL2RA regulators formed a regulatory connection with at least one, and often many, other IL2RA regulators. These data highlight that each regulator’s effect on a target gene is likely mediated by numerous other regulators. Furthermore, precise gene expression levels are also maintained through numerous feedback loops, which we could detect since we individually knocked out each regulator. As we only measured a subnetwork focused on the 24 genes that we knocked out, the full transcriptional network likely contains many additional complicated feedback loops. As technology continues to improve, we will likely need to scale to both genome-wide perturbations and genome-wide profiling to build comprehensive gene regulatory network maps.

Our strategy to map gene networks is likely broadly applicable to many situations where we know a few genes of interest but have yet to discover what other genes are involved. A landmark paper in yeast perturbed 1,484 regulatory genes and profiled the resulting transcriptional changes, demonstrating how large perturbation studies can be used to assign genes to functional pathways (Kemmeren et al., 2014). As similar approaches at this scale are not yet feasible in other systems, we used a two-step mapping approach. We first identified the most important regulatory genes and then performed a more focused set of perturbations coupled with RNA-Seq and ATAC-Seq. The strong enrichment for immune and immune disease genes among highly co-regulated genes suggest that this is a powerful and broadly applicable approach for identifying functionally related genes.

The identified IL2RA regulators co-regulate a central network highly enriched for genes that are associated with immune diseases. This network architecture is consistent with the omnigenic model that we proposed, in which a set of core genes act directly on a trait, but perturbations of many peripheral genes affect the expression of these core genes (Boyle et al., 2017; Liu et al., 2019). This network structure could explain how the dysregulation of many seemingly unrelated GWAS hits could disrupt a central network of important disease genes. Although the functional importance of such a co-regulated network is unclear, it will be interesting to determine whether a similar network structure exists in other cell types and whether targeting key regulators of this central network is an effective therapeutic strategy. Additionally, the generation of this network has great potential for the understanding of immune disease, including potential relevance for the diagnosis of the genetic causes of primary immunodeficiencies. In the future, candidate genetic variants found in clinical genome sequencing results that map to in this highly co-regulated immune network could be prioritized for follow-up, even if they have not previously been linked to immune dysregulation diseases. Lastly, IL2RA, IL-2, and CTLA4, which served as seed genes for this immune disease network, are established targets of drug development for cancer and autoimmune diseases (Mullard, 2021; Rowshanravan et al., 2018); this network will likely help to identify promising new drug targets.

Assembling gene network maps can also be used to better engineer immune cell therapies. T cells with defined T cell receptors or chimeric antigen receptors (CARs) that specifically target tumors are being employed in the clinic with incredible success (Esensten et al., 2017). Our group and others are developing methods to both disrupt and overexpress genes to further engineer T cells for immunotherapies (Lynn et al., 2019; Roth et al., 2020; Stadtmauer et al., 2020). By identifying regulators that control the expression of key immune genes and mapping the genome-wide networks they control, we have new opportunities to engineer T cell therapies with improved cell fitness, cytokine production, and cytotoxic function. As gene engineering moves into the clinic, understanding gene regulatory networks will allow for the safe manipulation of genes with a more holistic understanding of the downstream direct and indirect consequences of such manipulation.

## Supporting information

Table S1 sgRNA library

Table S2 screen gene results

Table S3 screen sgRNA results

Table S4 RNA ATAC sgRNAs

Table S5 RNA-Seq results

Table S6 ATAC-Seq results

Table S7 CRISPR genotyping primers

Table S8 sequencing primers

## Acknowledgements

We thank members of the Marson and Pritchard labs for helpful discussions and manuscript feedback. We thank Harold Pimentel for advice on analysis. This research was supported by NIH grants R01HG008140 and RM1-HG007735. A.M. holds a Career Award for Medical Scientists from the Burroughs Wellcome Fund, is an investigator at the Chan Zuckerberg Biohub and has received funding from the Innovative Genomics Institute (IGI), the American Endowment Foundation, the Cancer Research Institute (CRI), the Lloyd J. Old STAR award, a gift from the Jordan Family, a gift from Barbara Bakar, and is a member of the Parker Institute for Cancer Immunotherapy (PICI). O.S. was supported by the National Institutes of Health grant number T32AI125222. S.N. was supported by a Helen Hay Whitney Fellowship. N.S.-A was supported by a Stanford Graduate Fellowship and CEHG Fellowship. A.F.C. was supported by an NIH F32 postdoctoral fellowship (5F32GM135996-02). Sorting was carried out at the UCSF Flow Cytometry Core (RRID:SCR_018206) supported in part by Grant NIH P30 DK063720 and by the NIH S10 Instrumentation Grant S10 1S10OD021822-01. The RNA-Seq was carried out at the DNA Technologies and Expression Analysis Cores at the UC Davis Genome Center, supported by NIH Shared Instrumentation Grant 1S10OD010786-01. Other sequencing was carried out at the UCSF CAT, supported by a PBBR grant. Some of the computing for this project was performed on the Sherlock cluster. We would like to thank Stanford University and the Stanford Research Computing Center for providing computational resources and support that contributed to these research results.

## Author Contributions

Conceptualization, J.W.F, O.S., J.K.P, and A.M.;

Formal Analysis, J.W.F, S.N, and N.S.-A.;

Investigation, J.W.F., O.S., A.K., A.F.C. and J.C.;

Resources, W.J.G, J.K.P, and A.M.;

Writing - Original Draft, J.W.F;

Writing - Review & Editing, J.W.F, O.S., J.K.P, and A.M.;

Visualization, J.W.F;

Supervision, W.J.G, J.K.P, and A.M.;

Funding Acquisition, W.J.G, J.K.P, and A.M.;

## Declaration of Interests

A.M. is cofounder, member of the Boards of Directors and member of Scientific Advisory Boards of Spotlight Therapeutics and Arsenal Biosciences. A.M. has served as an advisor to Juno Therapeutics, was a member of the scientific advisory board at PACT Pharma and was an advisor to Trizell. A.M. has received honoraria from Merck and Vertex, a consulting fee from AlphaSights, and is an investor in and informal advisor to Offline Ventures. A.M. owns stock in Arsenal Biosciences, Spotlight Therapeutics and PACT Pharma. The Marson lab has received research support from Juno Therapeutics, Epinomics, Sanofi, GlaxoSmithKline, Gilead and Anthem. W.J.G. is a consultant for 10x Genomics, which has licensed IP associated with ATAC-seq. W.J.G. has additional affiliations with Guardant Health (consultant) and Protillion Biosciences (cofounder and consultant). J.W.F., O.S., J.K.P., and A. M. are listed as inventors on a patent application related to this work.

## Resource Availability

### Data and Code Availability

The raw sequencing files generated during this study are available at GEO: GSE171737.

## Methods

### Pooled sgRNA library construction

We selected transcription factors (TFs) with known or inferred motifs from Lambert et al. (Lambert et al., 2018), non-target controls from the Brunello sgRNA library (Doench et al., 2016), hits from a previous screen (Cortez et al., 2020) and immune genes of interest from the lab. All sgRNA sequences were taken from the Brunello sgRNA library (Doench et al., 2016). In total we included 1349 genes with an average of 4 guides per gene, 13 guides against GFP as a positive control for editing, and 593 non-targeting controls. We ordered a pooled oligo library from Twist Biosciences with flanking sequences for cloning into the plasmid backbone. Following the custom sgRNA library cloning protocol as described by Joung et al. (Joung et al., 2017), we integrated our sgRNA library into the LRG2.1 backbone featuring an improved guide RNA scaffold (Addgene, plasmid# 108098) from Grevet et al. (Grevet et al., 2018) using NEBuilder HiFi DNA Assembly master mix (NEB, Cat #E2621X) according to the manufacturer’s protocol. We used Endura ElectroCompetent Cells to amplify the library per the manufacturer’s protocol (Endura, Cat #60242-1). Finally we performed maxipreps using the ZymoPure Plasmid Maxiprep kit (Zymo, Cat # D4202).

### Lentiviral production

HEK 293T cells were seeded at 14 million cells in a 15 cm tissue culture treated culture dish (Corning, Cat #430599) in Opti-MEM (UCSF CCF, Cat #CCFAC008) 24 hours prior to transfection. Using Lipofectamine 3000 (Lifetech, Cat #L3000075) according to the manufacturer’s protocol, cells were transfected with the sgRNA transfer plasmid, and two lentiviral packaging plasmids, pMD2.G (Addgene, Cat #12259) and psPAX2 (Addgene, Cat #12260). Cells were incubated for 5 hours at 37°C, after which time the transfection media was replaced with fresh Opti-MEM containing ViralBoost at 1x (Alstem, Cat #VB100). Cells were incubated for 24 hours and then the viral supernatant was collected and spun down at 300*g* for 5 minutes to remove cellular debris. The supernatant was then passed through a 0.45-µm filter, and subsequently mixed with one volume of cold Lentivirus Precipitation Solution (Alstem, Cat #VC125) at 4° C to every 4 volumes of lentivirus-containing supernatant. Samples were mixed well and placed at 4°C overnight. The virus was then concentrated by centrifugation at 1500*g* for 30 minutes at 4°C, after which the supernatant was discarded, and the residual sample underwent additional centrifugation at 1500*g* for 5 minutes to remove any residual supernatant. The viral pellet was then resuspended at a ratio of 1:100 of the original volume using PBS (Fisher Scientific, Cat #10010049) at 4°C. Virus was then stored until use at −80°C.

### Isolation, culture and expansion of human CD4+ T-effector cells

Primary human T cells were obtained from residuals from leukoreduction chambers after apheresis (Blood Centers of the Pacific) for experiments not involving amplicon-, RNA-, or ATAC-sequencing. For sequencing experiments, primary human T cells were obtained from whole blood donors through a protocol approved by the UCSF Committee on Human Research (CHR#13-11950) or through consented Leukopaks (STEMCELL). Peripheral blood mononuclear cells (PBMCs) were isolated by size separation using Lymphoprep (STEMCELL, Cat #07861) in SepMate tubes (STEMCELL, Cat #85460), according to the manufacturer’s protocol. CD4+ T cells were isolated from the responder population using the StemCell EasySep Human Isolation Kit (Catalog # 18063). Isolated cells were stimulated with Immunocult Human CD3/CD28/CD2 T Cell Activator (STEMCELL, Cat #10970) at 6.25 uL per 1E6 cells, cultured in RPMI with 50 U/mL IL-2 (Amerisource Bergen, Cat #10101641) at a concentration of 1E6 cells/mL.

### Culture media

Cells were grown in RPMI (Sigma, Cat # R0883) with 10% FCS (Sigma, Cat # F0926), with 100U/mL Pen-Strep (Gibco, Cat # 15140-122), 2mM L-Glutamine (Sigma, Cat # G7513), 10mM HEPES (Sigma, Cat # H0887), 1X MEM Non-essential Amino Acids (Gibco, Cat # 11140-050), 1mM Sodium Pyruvate (Gibco, Cat # 11360-070), and 50 U/mL IL-2 (Amerisource Bergen, Cat #10101641) at a concentration of 1E6 cells/mL.

### Pooled CRISPR screen

#### Lentiviral transduction

Twenty-four hours post stimulation, lentivirus containing the sgRNA library was added directly to cultured T cells in a drop-wise fashion while tilting the flasks to distribute evenly, targeting a multiplicity of infection (MOI) of 0.4. After an additional 24 hours, excess lentivirus was removed from the supernatant and washed off the cells. Cells were then incubated at 37°C.

#### Cas9-ribonucleotide protein (RNP) preparation

Cas9 protein (MacroLab, Berkeley, 40 µM stock) was delivered into the cells using a modified Guide Swap technique (Ting et al., 2018). Lyophelized Dharmacon Edit-R crRNA Non-targeting Control #3 (Dharmacon, Cat #U-007503-01-05) and Dharmacon Edit-R CRISPR-Cas9 Synthetic tracrRNA (Dharmacon, Cat #U-002005-20) were resuspended at a stock concentration of 160 mM in 10 mM Tris-HCl (pH 7.4) with 150 mM KCl. They were mixed at a 1:1 ratio, creating an 80 mM solution, and incubated on a heat block at 37°C for 30 minutes. Single-stranded donor oligonucleotides (ssODN; sequence: TTAGCTCTGTTTACGTCCCAGCGGGCATGAGAGTAACAAGAGGGTGTGGTAATATTACGGTACCGAGCACTATCGATACAATATGTGTCATACGGACACG) was then added at a 1:1 molar ratio of the final Cas9-Guide complex, and mixed well by pipetting. The solution was incubated for an additional 5 minutes at 37°C on the heat block. Cas9 was then added slowly at a 1:1 volume to volume ratio, taking care to avoid precipitation, pipetting up and down several times to ensure complete resuspension of the RNP complex, and incubated at 37°C for 15 minutes.

#### Electroporation

24 hours after virus was washed from the culture, cells were centrifuged at 100*g* for 10 minutes to pellet them, and resuspended in room temperature Lonza P3 electroporation buffer (Lonza, Cat #V4XP-3032) at 1-2E6 cells per 17.8 µL. 7.2 µL of the RNP-ssODN complex were added for every 17.8 µL of cells and mixed well. Using a multichannel pipette, 23 uL of the cells-RNP-ssODN mixture were added to each well of a 96 well electroporation cuvette plate (Lonza, Cat #VVPA-1002), and nucleofected using the pulse code EH-115. Immediately after electroporation, 90 µL of prewarmed media were added to each well and incubated at 37°C for 15 minutes. Cells were then pooled, transferred to incubation flasks, and diluted with pre-warmed media to a final concentration of 1E6 cells/mL and incubated at 37°C. Cells were passaged at 48 hours post electroporation, and subsequently maintained in culture at 1E6 cells/mL.

### Screen phenotyping and cell sorting

Cells were collected for analysis 6 days after electroporation. 10-20E6 cells were portioned off and sorted based on GFP expression only. The remaining cells were sorted based on GFP positivity, as well as a target expression using an APC fluorescent antibody targeting either IL2RA (CD25) (Tonbo, Cat #20-0259-T100), IL-2 (Biolegend, Cat #500310), or CTLA-4 (Biolegend, Cat #349908). Cells sorted for IL2RA underwent surface staining according to the manufacturer’s protocol. Cells sorted for IL-2 were treated with Cell Activation Cocktail with Brefeldin A (Biolegend, Cat #423304) for 4 hours prior to fixation, and were fixed using the BD Cytofix/Cytoperm kit (Becton Dickinson, Cat #554714) according to the manufacturer’s protocol. Cells sorted for CTLA-4 were treated with Cell Activation Cocktail without Brefeldin A (Biolegend, Cat #423302) for 4 hours prior to fixation, and were fixed using the Foxp3 Fix/Perm buffer set (Biolegend, Cat #421403) according to the manufacturer’s protocol. Cells were sorted using a BD FACS Aria II. The IL2RA screen was performed in 3 donors, while the IL-2 and CTLA4 screens were performed in 2 donors.

### Genomic DNA extraction and preparation for next generation sequencing

After sorting, cells were washed with PBS, counted, pelleted, and resuspending at up to 5E6 cells per 400 µl of lysis buffer (1% SDS, 50 mM Tris, pH 8, 10 mM EDTA). The remaining protocol reflects additives/procedures performed per each 400 µl of sample. 16µl of NaCl (5M) was added, and the sample was incubated on a heat block overnight at 66°C. The next morning, 8µl of RNAse A (10mg/ml, resuspended in ddH-_2_O) (Zymo, Cat #E1008) was added, and the sample was vortexed briefly, and incubated at 37°C for 1 hour. Next, 8µl of Proteinase K (20mg/ml) (Zymo, Cat #D3001) was added, the sample was vortexed briefly, and incubated at 55°C for 1 hour. A phase lock tube (Quantabio, Cat #2302820) was prepared for each sample by spinning down the gel to the bottom of the tube at 20,000g for 1 minute and then 400µl of Phenol:Chloroform:Isoamyl Alcohol (25:24:1) was added to each tube. 400µl of the sample was then added to the phase lock tube and the tube was shaken vigorously. The sample was centrifuged at maximum speed at room temperature for 5 minutes. The aqueous phase was transferred to a low-binding eppendorf tube (Eppendorf, Cat #022431021) and then 40µl of Sodium Acetate (3M), 1µl GlycoBlue (Invitrogen, Cat # AM9515), and 600µl of room temperature isopropanol was added. The sample was then vortexed and stored at −80°C for 30 minutes or until the sample had frozen solid. Next the sample was centrifuged at maximum speed at 4°C for 30 minutes, the pellet was washed with fresh 70% room temperature Ethanol, and allowed to air dry for 15 minutes. Pellets were then resuspended in Zymo DNA elution buffer (Zymo, Cat No: D3004-4-10), and placed on the heat block at 65°C for 1 hour to completely dissolve the genomic DNA.

sgRNA was amplified and barcoded from the genomic DNA according to the protocol by Joung et al. (Joung et al., 2017). Up to 2.5 µg of genomic DNA were added to each 50 µL reaction, which included 25 µL of NEBNext Ultra II Q5 master mix (NEB, Cat #M0544L), 1.25 µL of the 10 µM forward primer and 1.25 µL of the 10 µM reverse primer, and H2O to 50 uL. The following PCR cycling conditions were used: 98°C for 3 minutes, followed by 23 cycles at 98°C for 10 seconds, 63°C for 10 seconds, and 72°C for 25 seconds, and ending with 2 minutes at 72°C. Samples were then cleaned and concentrated in Zymo Spin-V columns (Zymo, Cat #C1016-50) following Joung et al., and eluted in 150 uL of Zymo DNA elution buffer. Up to 2 µg of each library were loaded on a 2% agarose gel, and the band at ∼250 base pairs was extracted using the Zymoclean Gel DNA recovery kit (Zymo, Cat #D4008). The concentration of each sample was then measured using the Qubit dsDNA high sensitivity assay kit (Thermo Fisher Scientific, Cat #Q32854). Samples were then sequenced on an Illumina HiSeq 4000 using 10-30% PhiX (Illumina, Cat #15017872), and a custom sequencing primer. Primer sequences are listed in Table S8.

### Arrayed validation isolation, culture, and electroporation

We selected 50 genes, 3 positive controls (IL2RA, IL-2, and CTLA4), and 4 non-targeting controls for validation. Thirty-nine of these genes regulate at least two out of three target genes, while the remaining genes were selected because they had large effects in individual screens. For each gene, we selected the top two performing guides from the screen data. Primary human T cells were obtained from whole blood donors through a protocol approved by the UCSF Committee on Human Research (CHR#13-11950), isolated and stimulated as described above. Custom crRNA plates were ordered from Dharmacon, and were assembled as RNP-ssODN complexes as described above. 48 hours after stimulation, cells were counted, pelleted, and resuspended in room temperature Lonza P3 buffer (Lonza, Cat #V4XP-3032) at 1E6 cells per 20 µL. Cells were then mixed with 100 pmol of RNP, transferred to a 96 well electroporation cuvette plate (Lonza, Cat #VVPA-1002), and nucleofected using the pulse code EH-115. After electroporation, 90 µL of pre-warmed media was immediately added to each well and plates were incubated at 37°C for 15 minutes. Wells were then split to a target culture population of 1E6 cells/mL filling all edge wells in the 96-well plate with PBS in order to avoid edge-effects and incubated at 37°C.

### Arrayed validation phenotyping using flow cytometry and genotyping

Arrayed validation plates were phenotyped at 5 days after electroporation using the sample protocol and materials as outlined in the screen in a 96-well plate format. Cells were stained for IL2RA (CD25) (Tonbo, Cat #20-0259-T100), IL-2 (Biolegend, Cat #500310), or CTLA-4 (Biolegend, Cat #349908) and analyzed with an Attune NxT Flow Cytometer with a 96-well plate-reader.

On day 5 (sgRNA #1 Donor 1-3) or day 7 (sgRNA #2 Donor 1, 3) post-electroporation, genomic DNA was isolated from each sample using DNA QuickExtract (Lucigen, Cat #QE09050) according to the manufacturer’s protocol. Primers were designed to flank each sgRNA genomic target site. Amplicons containing CRISPR edit sites were generated by adding 1.25 µL each of forward and reverse primer at 10nM to 5 µL of sample in QuickExtract, 12.5 µL of NEBNext Ultra II Q5 master mix (NEB, Cat #M0544L), and H2O to a total 25 µL reaction volume. The following PCR cycling conditions were used: 98°C for 3 minutes, 15 cycles of 94°C for 20 seconds followed by 65°C-57.5°C for 20 seconds (0.5°C incremental decreases per cycle), and 72°C for 1 minute, and a subsequent 20 cycles at 94°C for 20 seconds, 58°C for 20 seconds and 72°C for 1 minute, and a final 10 minute extension at 72°C. For some samples with non-specific amplification, the above PCR was repeated with a higher temperature touchdown annealing step: 70°C-64°C (0.5°C incremental decreases per cycle). Samples were then diluted 1:200 and subsequently Illumina sequencing adapters and indices were added in a second PCR reaction. Indexing reactions included 1 µL of the diluted PCR1 sample, 2.5 µL of each the forward and reverse indexing primers at 10 µM each, 12.5 µL of NEB Q5 master mix, and H2O to a total 25 µL reaction volume. The following PCR cycling conditions were used: 98°C for 30 seconds, followed by 98°C for 10 seconds, 60°C for 30 seconds, and 72°C for 30 seconds for 12 cycles, and a final extension period at 72°C for 2 minutes. Samples were quantified in a 96-well plate reader using the Quant-IT DNA high sensitivity assay kit (Invitrogen, Cat #Q33232) according to the manufacturer’s protocol and pooled into 1 tube. Post pooling, samples were then SPRI purified, and quantified using an Agilent 4200 TapeStation. Samples were then sequenced on an Illumina MiniSeq or NextSeq with PE 300 reads. Primer sequences listed in Tables S7 and S8.

### Collection for RNA-Seq and ATAC-Seq

The top 24 hits with the largest effects in the IL2RA validation were selected for genomic profiling via RNA-Seq and ATAC-Seq. T cells were isolated from 3 donors using consented Leukopaks (STEMCELL) as described above. For each gene we used the sgRNA sequence that had the largest effect in the validation and ordered custom crRNAs from Dharmacon. We also ordered 8 custom crRNAs targeting the safe harbor AAVS1 locus (AAVS1 #1-8). These crRNAs were assembled as RNP-ssODN complexes and electroporated into cells as described above. Five days after electroporation, 60,000 cells were collected and used to generate ATAC-Seq libraries. A fraction of each sample was used for IL2RA flow cytometry as described above to obtain matched protein and RNA expression changes. The remaining cells were lysed with Zymo QuickRNA lysis buffer and isolated with the Zymo QuickRNA micro kit (Zymo, Cat #R1050) using the manufacturer’s protocol. RNA samples were treated with 1.5 ul of Turbo DNase (Invitrogen, Cat # AM2238) and then cleaned up using the Zymo RNA-5 Clean and Concentrator (Zymo, Cat #R1016). Isolated RNA integrity and concentration was checked using Agilent RNA Screen Tapes (Agilent, Cat #5067-5576)

### RNA-Seq

RNA was submitted to the UC Davis DNA Technologies and Expression Analysis Core to generate 3’ Tag-Seq libraries with unique molecular indices (UMIs). Barcoded sequencing libraries were prepared using the QuantSeq FWD kit (Lexogen, Vienna, Austria) for multiplexed sequencing according to the recommendations of the manufacturer (Lexogen). The fragment size distribution of the libraries was verified via micro-capillary gel electrophoresis on a Bioanalyzer 2100 (Agilent, Santa Clara, CA). The libraries were quantified by fluorometry on a Qubit fluorometer (LifeTechnologies, Carlsbad, CA), and pooled in equimolar ratios. Samples were sequenced on a HiSeq 4000 sequencer (Illumina, San Diego, CA).

### Plate-ATAC protocol

We harvested, counted, and treated each T cell culture with 200U/mL of DNase (Worthington Cat # LS002007) for 30 mins at 37 degrees. We then transferred 60,000 cells of each T cell culture into individual wells of a 96-well plate and washed cells once with PBS and once with RSB (10mM Tris HCl pH 7.5, 10mM NaCl, 3mM MgCl2). Cells were lysed in 50uL of cell lysis buffer (0.1% NP40, 0.1% Tween-20, 0.01% Digitonin in RSB) on ice for 3 min. We then added 150ul of RSB with 0.1% Tween-20 to each well and pelleted nuclei at 500g for 10 min at 4 degree. Cells were resuspended in 50ul transposition mix (25ul 2X TD buffer, 16.5ul 1X PBS, 0.5ul 10% Tween, 0.5ul 1% Digitonin, 2.5ul Tn5 transposase, and 5ul H2O) and transposition was performed at 37 degrees for 30 min with 300rpm shaking. Transposed fragments were purified using ZR-96 DNA Clean & Concentrator-5 Kit (Zymo D4024) and libraries were generated using PCR amplification with Nextera adapters and purified using Ampure beads. ATAC-Seq libraries were sequenced on a Novaseq with paired end 100 bp reads. The following samples produced low yield libraries and were not carried forward with sequencing: Donor 1 PTEN KO, Donor 3 STAT5B KO, Donor 3 AAVS1_4 control.

## Quantification and Statistical Analysis

### Analysis of pooled screens

A table of individual guide abundance in each sample was generated using the count command in MAGeCK version 0.5.8 (Li et al., 2014). Two individual guide RNAs (s_991 and s_3329) were filtered out due to extreme outlier counts. The MAGeCK test command was used to identify differentially enriched sgRNA targets between the low and high bins. All genes with an FDR adjusted p-value < 0.05 were considered significant. For screen QC, classification of essential genes, fitness, and non-essential genes were taken from Hart et al. (Hart and Moffat, 2016). Screen targets were classified as expressed if they had a count greater than 0 transcripts per million in CD4+ T cell RNA-Seq from (Calderon et al., 2019).

### Analysis of arrayed validation

Cells were gated on Lymphocytes and singlets in FlowJo Version 10.1 and the median fluorescence intensity (MFI) for APC-height for each stain was exported to csv files. Fluorescence data was imported into R. The MFI across 4 non-targeting controls was averaged per donor per plate and the MFI of each well on the plate was then normalized to the average MFI of the control wells. Individual wells with less than 30% lymphocytes were excluded from analysis due to toxicity. Knockout of HINFP appeared to be toxic and caused most of the cells to die so was excluded from analysis.

### Analysis of RNA-Seq data

Adapters were trimmed from fastq files using cutadapt version 2.10 (Martin, 2011) with default settings keeping a minimum read length of 20 bp. Reads were mapped to the human genome GRCh38 keeping only uniquely mapping reads using STAR version 2.7.b (Dobin et al., 2013) with the following settings “--outFilterMultimapNmax 1”. UMIs were extracted from fastqs using umi_tools version 1.0.1 (Smith et al., 2017) extract command with the following settings “--extract-method=regex --bc-pattern=’(?P<UMI_1>.{{6}})(?P<DISCARD_1>.{{4}}).*’”. Reads were then deduplicated using umi_tools dedup command with default settings. Deduplicated reads overlapping genes were then counted using featureCounts version 2.0.1 (Liao et al., 2014) with the following settings “-s 1” and using the Gencode version 35 basic transcriptome annotation.

To identify differentially expressed genes, UMI deduplicated counts between each KO sample and AAVS1 control samples were compared; sample Donor 4 AAVS1 #6 was excluded as an outlier. A minimum count per million was calculated based on the read depth of the samples being compared using the following command “ceiling(10/(min(colSums(count_mat))/1e6))” in R; genes with less reads than this minimum count across at least 3 samples were filtered out. Significantly differentially expressed genes for each KO were then identified using limma version 3.44.3 (Ritchie et al., 2015) while controlling for any differences between donors. Significant differentially expressed genes were defined as having a FDR adjusted p-value < 0.05.

### Analysis of ATAC-Seq data

Adapters were trimmed from fastq files using cutadapt version 2.10 (Martin, 2011) with default settings keeping a minimum read length of 20 bp. Reads were then mapped to the human genome GRCh38 using bowtie2 version 2.4.1 (Langmead and Salzberg, 2012) with the following settings “-X 2000 --very-sensitive”. Low quality reads were filtered using samtools version 1.10 (Li et al., 2009) using the following command “samtools view -h -b -F 1804 -f 2 -q 30”. Reads mapping within the ENCODE blacklist region were removed using bedtools version 2.29.2 (Quinlan and Hall, 2010) using bedtools intersect. Duplicated reads were removed using picard version 2.23.3 (http://broadinstitute.github.io/picard/) using the following settings “VALIDATION_STRINGENCY=LENIENT”. Reads mapping to ChrX, ChrY, and ChrM were excluded from further analysis. Reads were converted to single nucleotide ATAC insertion sites using the following command “bedtools bamtobed -i {input.bam} | awk ‘BEGIN {{OFS = “\t”}} $6 == “+” {{$2 = $2 + 4; $3 = $2 + 1; print}} $6 == “-&” {{$3 = $3 -4; $2 = $3 - 1; print}}’ | sort -k1,1 -k2,2n > {output.bed}”. For each sample, peaks were called using MACS2 version 2.2.7.1 (Zhang et al., 2008) using the ATAC insertion site bed files as input with the following settings “--format BED --shift -75 --extsize 150 -- nomodel --call-summits --nolambda --keep-dup all -B --SPMR -q 0.01”.

Peaks called in individual samples were merged into a consensus peak file in the following way using single nucleotide peak summits from MACS2. For each KO or control condition, peak summits that were within 75 bp of another peak summit in two out of three donors and were supported by at least 10 ATAC-Seq reads were merged into reproducible summit clusters. Across all samples, reproducible summits from each KO or controls were aggregated with other summits within 150 bp of each other.

For each aggregate cluster, we calculated an average summit location based on the location of all of the summits within the cluster. Each average summit location was then extended to 350 bp peaks centered on the average summit location to generate a consensus peak list. For each sample, the number of ATAC-Seq insertion sites that overlap each consensus peak was counted using the summarizeOverlaps function in the GenomicAlignments package (Lawrence et al., 2013).

To identify differentially accessible peaks, counts between each KO sample and all AAVS1 control samples were compared. A minimum count per million was calculated based on the read depth of the samples being compared using the following command “ceiling(10/(min(colSums(count_mat))/1e6))” in R; peaks with less reads than this minimum across at least 3 samples were filtered out. Significantly differentially accessible peaks for each KO were then identified using DESeq2 version 1.28.1 (Love et al., 2014) while controlling for both donor and the enrichment of reads at the TSS, which controls for the sample quality of individual ATAC-Seq samples. Significant differentially accessible peaks were defined as having a FDR adjusted p-value < 0.05.

### Motif enrichment in ATAC-Seq peaks

Known transcription factor binding motifs were downloaded from CIS-BP (Weirauch et al., 2014), JASPAR2020 (Fornes et al., 2020) and HOCOMOCO (Kulakovskiy et al., 2018). Motifs in ATAC-Seq peaks were identified with the motifmatchr package version 1.10.0 (DOI: 10.18129/B9.bioc.motifmatchr) using the matchMotifs function with default settings. If a given transcription had multiple binding motifs listed across these three databases, any associated motif that fell within an ATAC-Seq peak was considered a match.

### Analysis of ChIP-Seq data

Preproccessed ChIP-Seq coverage bigwigs for GATA3 and ETS1 binding in CD4+ T cells were downloaded from ChIP-Atlas (Oki et al., 2018). ChIP-Atlas maps ChIP-Seq data to the human genome GRCh38 using Bowtie2, removes PCR duplicates with SAMtools, and calculates coverage in Reads Per Million mapped reads using bedtools. ETS1 ChIP-Seq data was sourced from Schmidl et al. (Schmidl et al., 2014) and GATA3 ChIP-Seq data from Kanhere et al. (Kanhere et al., 2012).

### Analysis of arrayed genotyping data

Adapter sequences were trimmed from fastq files using cutadapt version 2.8 (Martin, 2011) using default settings keeping a minimum read length of 50 bp. Insertions and deletions at each CRISPR target site were then calculated using Crispresso2 version 2.0.42 (Clement et al., 2019) with the following options “--quantification_window_size 3” and “--ignore_substitutions”. For each guide RNA, the calculated insertion/deletion frequencies were averaged between donors.

### Enrichment of immune genes, Mendelian disease genes, and autoimmune GWAS genes

Genes were associated with the gene ontology term “immune system process” (GO:0002376) using the bioconductor package org.Hs.eg.db version 3.11.4 (Carlson, 2017). A list of inborn errors of immunity Mendelian disease genes were downloaded from the International Union of Immunological Societies website (https://iuis.org/committees/iei/) December 2019 dataset or taken from the 2021 update (Tangye et al., 2021). Genome wide significant SNPs (p-value < 5*10^-8) associated with 24 autoimmune diseases were taken from Taylor et al (Taylor et al., 2021). Genes were categorized as autoimmune GWAS genes if their TSS is within 100kb of one of these SNPs. The average gene expression level for all expressed genes was calculated as the log2 average count per million across control AAVS1 #1, 2, 3, 4, 5, 7, 8 samples across all three donors. AAVS1 #6 was excluded due to one outlier sample. The significance of the association between these three gene categories and how highly a gene is co-regulated was calculated with the glm function in R using co-regulation bin and average gene expression as inputs.

### S-LDSC analysis

GWAS summary statistics were downloaded from the Price lab website (https://alkesgroup.broadinstitute.org/sumstats_formatted/ and https://alkesgroup.broadinstitute.org/UKBB/). LD scores were created for each annotation (corresponding to a set of differential ATAC-seq peaks or SNPs within 100kb of genes or their corresponding matched background sets) using the 1000G Phase 3 population reference. Each annotation’s heritability enrichment for a given trait was computed by adding the annotation to the baselineLD model and regressing against trait chi-squared statistics using HapMap3 SNPs with the stratified LD score regression package (Finucane et al., 2015). Heritability enrichments were meta-analyzed across immune or non-immune traits using inverse variance weighting. The ATAC-Seq background set was generated by randomly sampling peaks from all unchanged peaks. The ATAC-Seq peaks in the background set were matched to significant differential ATAC-Seq peaks based on deciles of chromatin accessibility in AAVS1 control cells. ATAC-Seq background peaks in each accessibility decile were further matched to differential peaks based on the percentage of proximal peaks (defined as within 2kb of a TSS). For each co-regulation bin, RNA-Seq background genes were sampled from the set of genes that were differentially expressed in less than 5 samples. RNA-Seq background genes were matched to significant differential RNA-Seq genes in each bin based on deciles of gene expression in AAVS1 control cells.

### Multiple sclerosis SNPs analysis

Multiple sclerosis SNPs from a recent multiple sclerosis GWAS meta-analysis (International Multiple Sclerosis Genetics Consortium, 2019) with pre-calculated Probabilistic Identification of Causal (PICS) scores were downloaded from pics2.ucsf.edu (Taylor et al., 2021). SNPs were filtered to be genome wide significant (p-value < 5*10^-8) and have a PICS probability > 0.5. SNPs that fall within 350 bp ATAC-Seq peaks were identified using the findOverlaps function in the GenomicRanges package version 1.40.0 (Lawrence et al., 2013).

### CTLA4, IL-2, and IL2RA disease associations

Clinical phenotypes associated with coding variants in CTLA4 and IL2RA were based on the International Union of Immunological Societies’ phenotype classifications (Bousfiha et al., 2020). Associations between CTLA4, IL-2, and IL2RA and common autoimmune diseases were based on connections between autoimmune GWAS SNPs and these genes on the Open Targets Platform (Ochoa et al., 2021).

### Plots and genomic tracks

All plots were generated in ggplot2 in R version 4.0.2 (Wickham, 2009). Network connections were visualized with ggraph package version 2.0.4 in R (https://CRAN.R-project.org/package=ggraph). Heatmap of ATAC-Seq changes at the IL2RA locus was visualized with Gviz version 1.32.0 in R (Hahne and Ivanek, 2016). For all ATAC-Seq coverage tracks, ATAC-Seq reads were extended to 100 bp centered on the ATAC-Seq insertion site. Size factors for normalization for each sample were estimated using the number of ATAC-Seq reads that fall in peaks with the estimateSizeFactorsForMatrix function from the DESeq2 package in R (Love et al., 2014). Reads at a particular genomic locus were then normalized by that sample’s size factor. Per base read coverage was averaged across all three donors and exported as a bigwig file. Of AAVS1 controls #1-8, AAVS1 control #4 was excluded from coverage calculations due to an insufficient number of reads sequenced across all three donors. ATAC-Seq and ChiP-Seq coverage at a particular locus was visualized using the pygenometracks package version 3.6 (Ramírez et al., 2018) with 25bp bins.

## Supplemental Figures

**Supplemental Figure 2:**
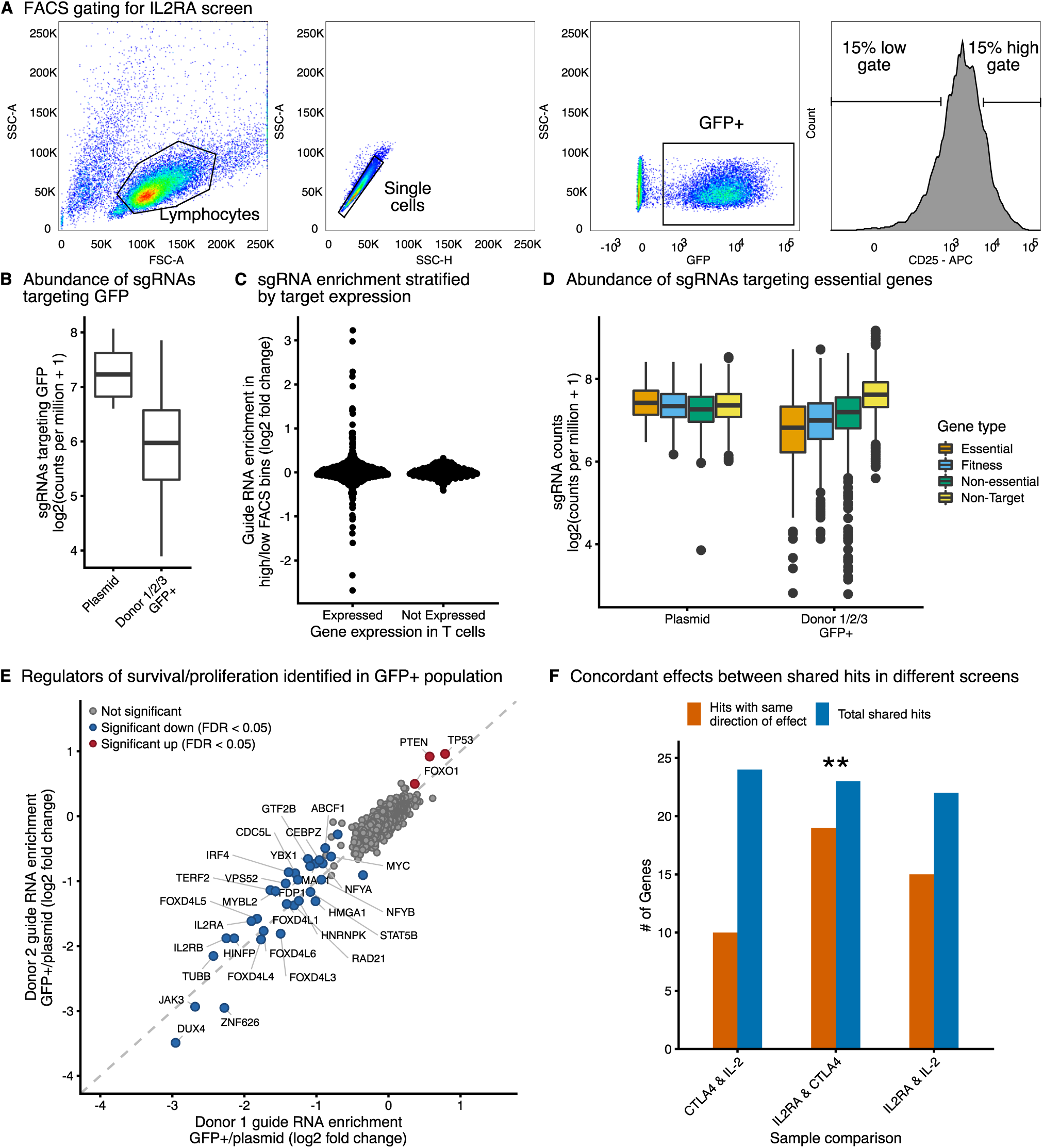
Quality control of the CRISPR screens. **A)** Fluorescence activated cell sorting gating for IL2RA, IL-2, and CTLA4 screens. Representative example from the IL2RA screen is shown. **B)** Abundance of guide RNAs targeting GFP in either the starting plasmid or in the GFP+ sorted population. **C)** Differential enrichment between the high-and low-expression bins for guide RNAs targeting genes that are either expressed or not expressed in CD4+ T cells based on RNA-Seq. **D)** Abundance of guide RNAs targeting essential genes, fitness genes, non-essential genes, or non-targeting guides in the starting plasmid or in the GFP+ sorted population in the three donors. **E)** Enrichment of guide RNAs between the GFP+ sorted population and starting plasmid. Results from Donor 1 and Donor 2 are depicted. Significant hits with an FDR adjusted p-value < 0.05 across all donors are highlighted. **F)** Comparison of the number of shared significant hits between the different screens and whether those hits have the same direction of effect on their targets. P-value < 0.01 sign test.

**Supplemental Figure 3:**
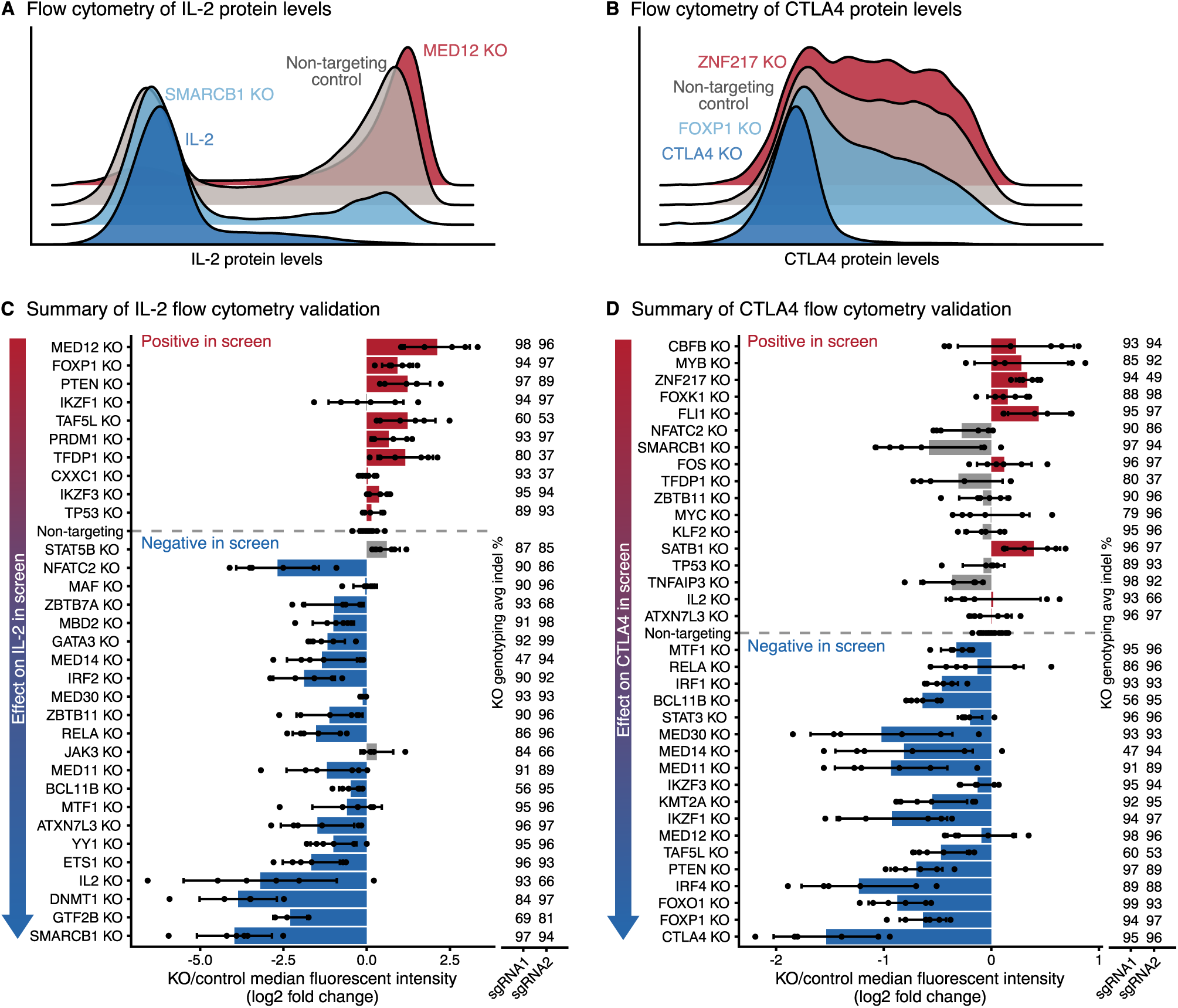
Arrayed KOs validate IL-2 and CTLA4 screen results. **A-B)** Representative flow cytometry density plots for IL-2 (A) or CTLA4 (B) protein levels after KO of top screen hits. KO of hits that decrease target levels are shown in blue and KO of hits that increase target levels are shown in red. **C-D)** Summary of changes in IL-2 (C) or CTLA4 levels measured using flow cytometry. Screen hits selected for validation are displayed on the Y-axis ordered by their effect size in the pooled CRISPR screen. For each KO, bars show the average change in IL-2 or CTLA4 median fluorescence intensity relative to non-targeting controls. Dots show individual data points and error bars show standard deviation across two guide RNAs and three donors per guide RNA. Concordant changes between the screen and validation that increase or decrease IL-2/CTLA4 levels are shown in red or blue respectively. Discordant changes are shown in grey. The average insertion/deletion (indel) percentage at the genomic target site across multiple donors for guide RNA 1 (n = *3)* and guide RNA 2 (n = 2) is shown to the right.

**Supplemental Figure 4:**
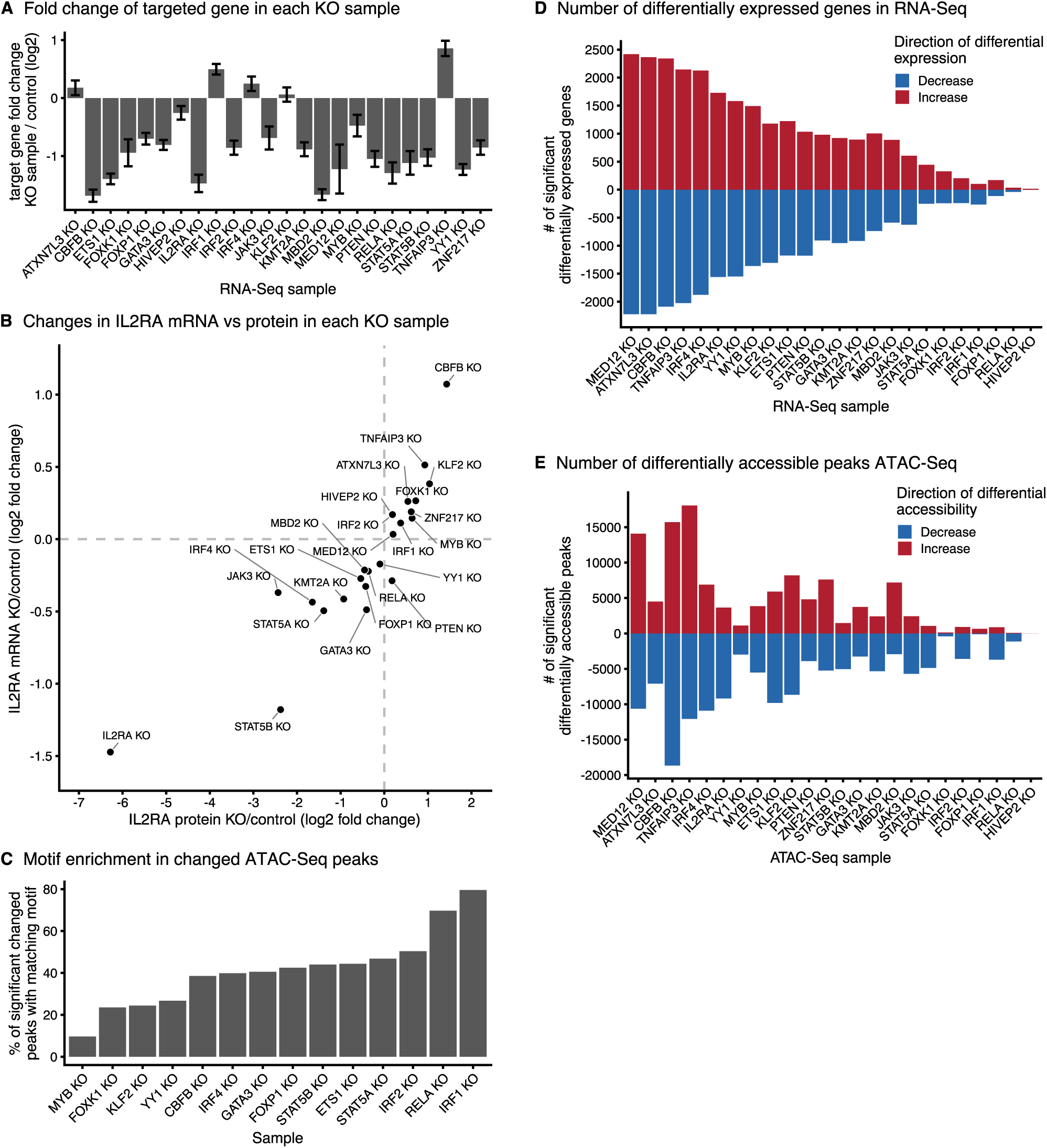
Mapping genome-wide transcripts and chromatin sites controlled by each IL2RA regulator. **A)** mRNA fold change for the CRISPR targeted gene in each KO sample. **B)** Comparison of average changes in IL2RA mRNA levels (RNA-Seq) and protein levels (flow cytometry) for each KO sample collected for RNA-Seq and ATAC-Seq. **C)** Percent of significantly changed ATAC-Seq peaks in each KO sample that contain a known motif for the knocked out transcription factor. **D-E)** The total number of significantly changed genes (D) or peaks (E) detected via RNA-Seq and ATAC-Seq in each KO sample.

**Supplemental Figure 6:**
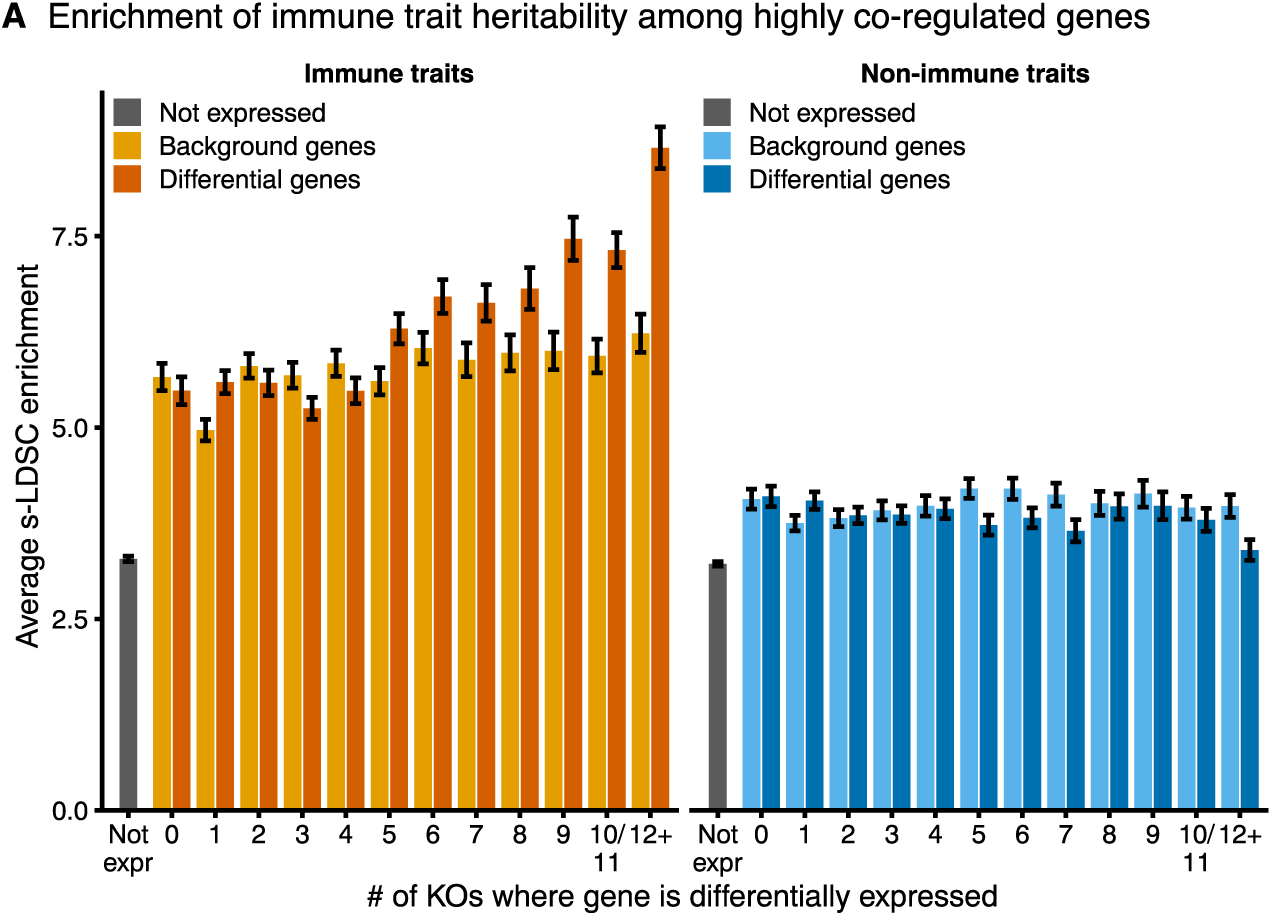
Highly co-regulated gene sets are enriched for immune disease genes. **A)** Enrichment of heritability for immune traits compared to non-immune traits in a window of 100kb around highly co-regulated genes. Enrichment for matched background gene sets for each co-regulation bin are shown. Enrichment calculated using stratified LD score regression. Traits were meta-analyzed using inverse-variance weighting; average enrichment and standard deviation shown.

**Supplemental Figure 7:**
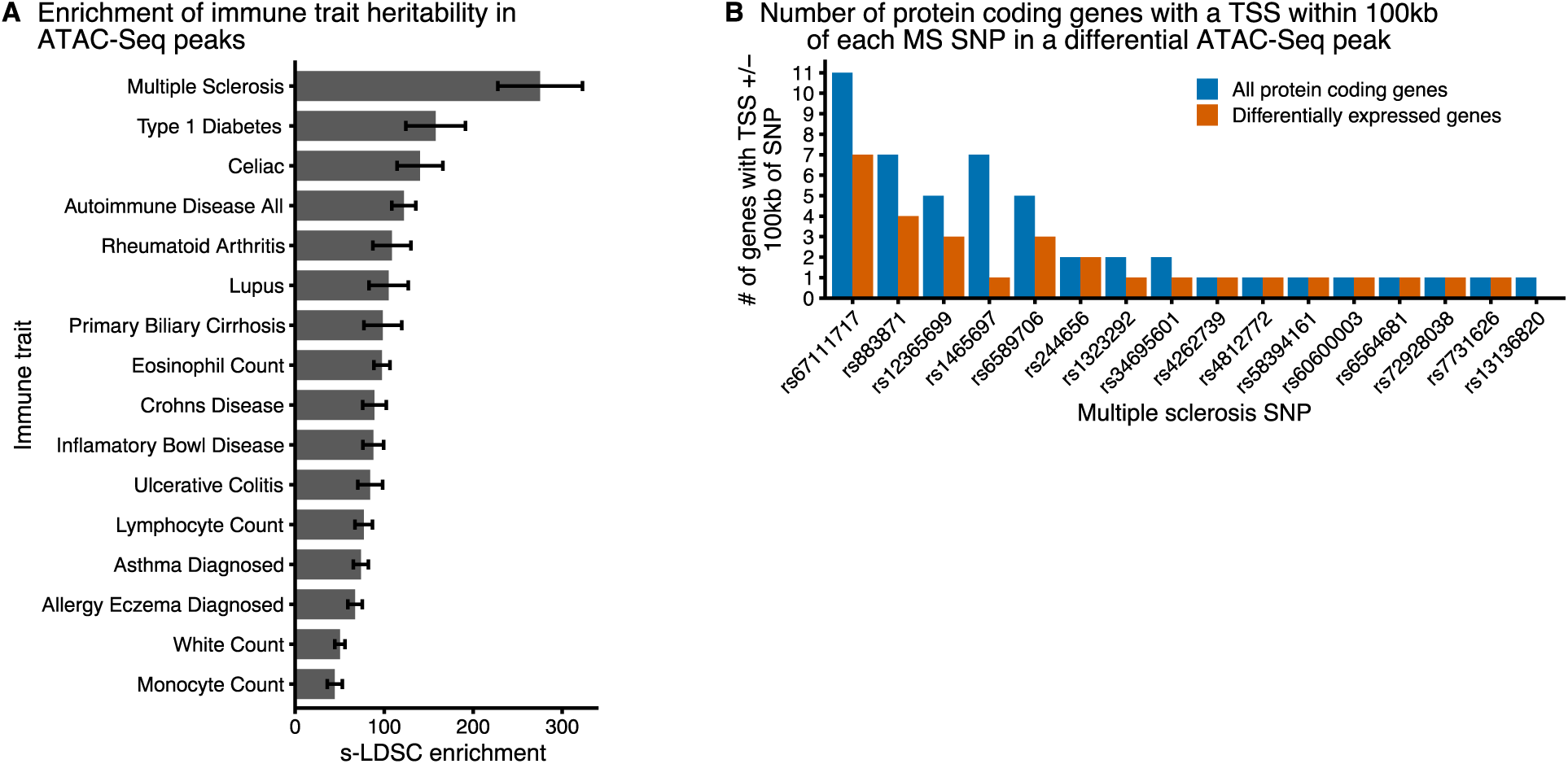
Multiple sclerosis (MS) SNPs within CD4+ T cell ATAC-Seq peaks. **A)** Enrichment of heritability in accessible ATAC-Seq peaks for different immune traits. Enrichment calculated with stratified LD score regression. **B)** The number of all protein coding genes and differentially expressed protein coding genes with a TSS within 100kb of a multiple sclerosis SNP. Only high confidence multiple sclerosis SNP within differentially accessible ATAC-Seq peaks are shown.

